# Communication between distinct subunit interfaces of the cohesin complex promotes its topological entrapment of DNA

**DOI:** 10.1101/559492

**Authors:** Vincent Guacci, Fiona Chatterjee, Brett Robison, Douglas Koshland

**Affiliations:** Department of Molecular and Cell Biology, UC-Berkeley, Berkeley, CA 94720

## Abstract

Cohesin mediates higher-order chromosome structure. Its biological activities require topological entrapment of DNA within a lumen(s) formed by cohesin subunits. The reversible dissociation of cohesin’s Smc3p and Mcd1p subunits are postulated to form a regulated gate that allows DNA entry and exit into the lumen. We assessed gate-independent functions of this interface in yeast using a fusion protein that joins Smc3p to Mcd1p. We show that in vivo all the regulators of cohesin promote DNA binding of cohesion by mechanisms independent of opening this gate. Furthermore, we show that this interface has a gate-independent activity essential for cohesin to bind chromosomes. We propose this interface regulates DNA entrapment by controlling the opening and closing of one or more distal interfaces formed by cohesin subunits, likely by inducing a conformation change in cohesin. Furthermore, cohesin regulators modulate the interface to control both DNA entrapment and cohesin functions after DNA binding.

## INTRODUCTION

Cohesin is a member of a family of Smc complexes required to higher order chromosome structure and dynamics (Onn *et al.*, 2008). The chromosomal binding of cohesin at multiple sites, termed CARs, serves to tether sister chromatids together from their formation in S phase through metaphase regions (Blat and Kleckner, 1999; Megee *et al.*, 1999; Laloraya *et al.*, 2000). This sister chromatid cohesion ensures proper chromosome segregation (Guacci *et al.*, 1997; Michaelis *et al.*, 1997). Cohesin is also important for mitotic chromosome condensation and proper gene expression (Guacci *et al.*, 1997; Rollins *et al.*, 2004). These functions of cohesin may arise from its ability to template chromosomes into distinct segments that could be packaged into domains of condensation and transcription (Guacci *et al.*, 1997; Hartman *et al.*, 2000; Lavoie *et al.*, 2002; Eagen, 2018). Finally, cohesin is loaded at sites of DNA breaks to facilitate both cohesion and repair (Strom *et al.*, 2007; Unal *et al.*, 2007). Understanding how cohesin carries out all these biological functions requires elucidating how cohesin binds to DNA, how binding is maintained and how binding is abrogated.

Cohesin must stably bind DNA to execute its different functions. In budding yeast the four evolutionarily conserved subunits of cohesin are Smc1p, Smc3p, Mcd1p/Scc1p and Scc3p (Guacci *et al.*, 1997; Michaelis *et al.*, 1997; Losada *et al.*, 1998). Biochemical studies indicate that DNA is topologically entrapped within cohesin, and such entrapment can occur within a trimer formed by Smc1p, Smc3p and Mcd1p (Ivanov and Nasmyth, 2007; Haering *et al.*, 2008; Gligoris *et al.*, 2014). The trimer architecture suggests it forms two lumens capable of trapping DNA. A large lumen is formed by dimerization of the Smc1p and Smc3p hinge domains at one end and dimerization of the two head domains at the other (Figure 1A). A smaller lumen is formed by the binding of the Mcd1p C-terminal domain to the Smc1p head domain, and the binding of the Mcd1 N-terminal helical domain (NHD) to coiled coil residues emerging from the Smc3p head domain (Figure 1B) (Haering *et al.*, 2008; Gligoris *et al.*, 2014). While initial models for topological entrapment favored DNA in the large lumen, studies of cohesin and other related SMC complexes have suggested entrapment by the small lumen as well (Stigler *et al.*, 2016).

**Figure 1:**
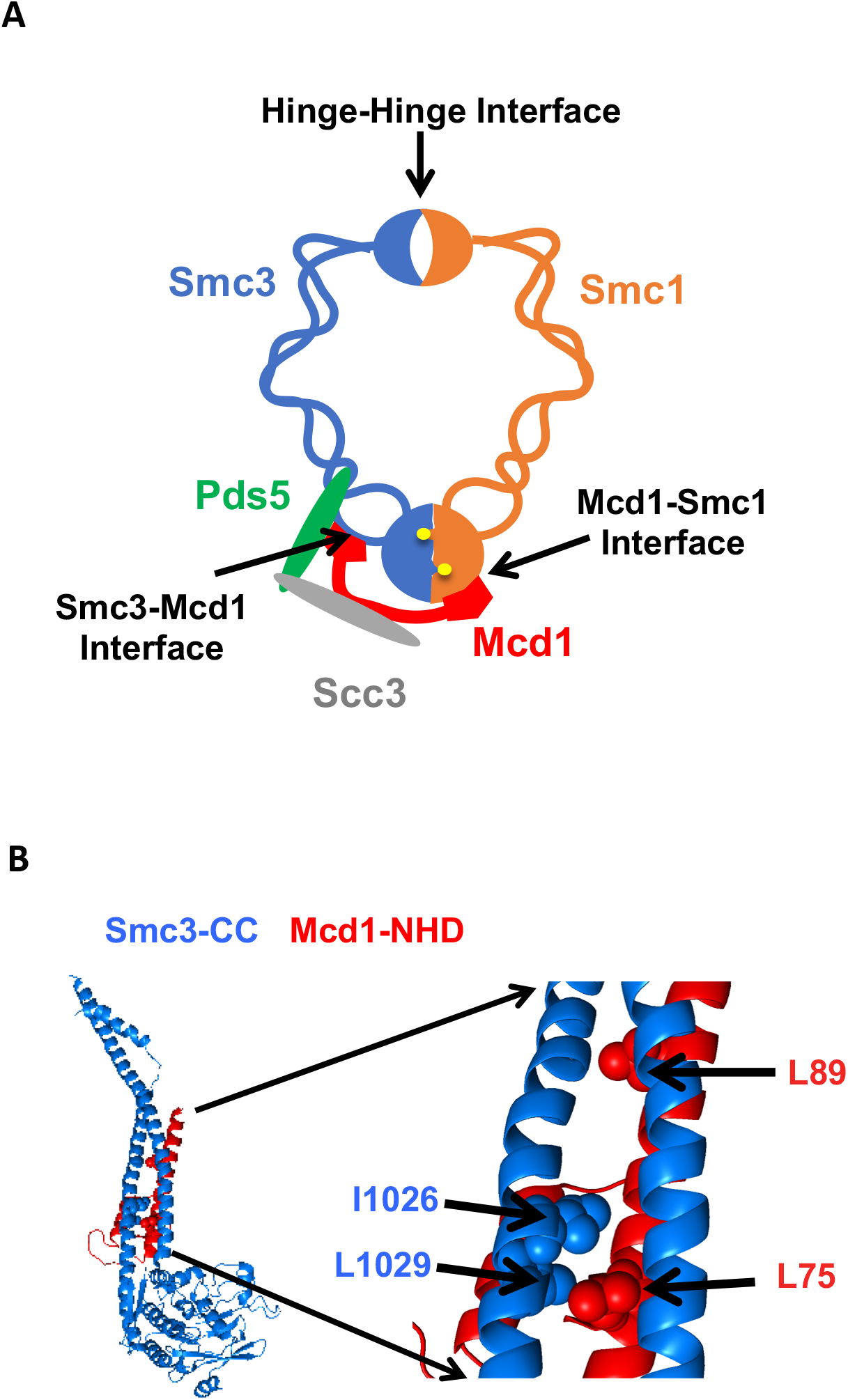
Cohesin structure. (A) Cartoon showing the cohesin complex and Pds5p. The three interfaces that can be chemically crosslinked to trap DNA within the Smc1p-Smc3p-Mcd1p trimer are marked by arrows. These are 1) Smc3p Mcd1p interface, 2) Smc1p hinge Smc3p hinge dimer interface, and 3) Smc1p Mcd1p interface. (B) Crystal structure of the Smc3p coiled-coil and Mcd1p NHD domain interface. Crystal structure entry PDB 4UX3 of the Smc3 head domain + short coiled-coil (Smc3 CC; blue) and Mcd1/Scc1 NHD (Mcd1 N; red) is shown. Right Side is enlargement showing essential residues in the interface as spheres.

To stably entrap DNA, one or more interfaces of the cohesin trimer must open to allow DNA to enter one of these lumens (Figure 1A). Controlling the opening and closing of these interfaces provides a mechanism for regulators to promote cohesin loading onto DNA, stable binding to DNA, and release from DNA. A dimeric Scc2p/Scc4p complex termed the cohesin loader is required for cohesion to bind DNA (Ciosk *et al.*, 2000). Evidence from *in vivo* cross-linking experiment indicate that DNA enters through opening of the hinge dimer interface and not through interfaces formed by head dimerization or interfaces between Smc3p and Mcd1p or Mcd1p and Smc1p (Gruber *et al.*, 2006). However, *in vitro* biochemical experiments suggested that Scc2/Scc4 may load cohesin onto DNA by the opening of the interface between Smc3p and Mcd1p (Murayama and Uhlmann, 2015). In this context, the Smc3p Mcd1p interface is a potential entry gate for topological entrapment.

Once loaded onto DNA, cohesin can either remain stably bound to it or be released. Toggling between these two states has been postulated to result from the antagonistic activities of the cohesin regulators, Eco1p, Pds5p and Wpl1p acting on or near the Smc3p Mcd1p interface. Eco1p (Ctf7p) mediated acetylation of the Smc3p K112 K113 head domain residues promotes stable cohesin DNA binding and cohesion (Skibbens *et al.*, 1999; Tóth *et al.*, 1999; Rolef Ben-Shahar *et al.*, 2008; Unal *et al.*, 2008; Bloom *et al.*, 2018). Once cohesion is established, Pds5p promotes cohesion maintenance as well as condensation (Hartman *et al.*, 2000; Stead *et al.*, 2003; Noble *et al.*, 2006; Robison *et al.*, 2018). Wpl1p binds to Pds5p to promote cohesin’s release from DNA and serves to antagonize both cohesion and condensation (Gandhi *et al.*, 2006; Kueng *et al.*, 2006; Rowland *et al.*, 2009; Guacci and Koshland, 2012; Lopez-Serra *et al.*, 2013; Bloom *et al.*, 2018)

Additional studies of Eco1p, Wpl1p and Pds5p have suggested a molecular basis for their ability to toggle cohesin binding to DNA. The binding of an Mcd1p N-terminal cleavage fragment to Smc3p is destabilized by Wpl1p, but is stabilized by Smc3p acetylation or by mutants that bypass *eco1* function (Chan *et al.*, 2012; Beckouet *et al.*, 2016). These experiments led to an exit gate model whereby DNA can escape cohesin entrapment by Wpl1p-dependent opening the Smc3p Mcd1p interface. Smc3p acetylation would inhibit Wpl1p, thereby keeping the interface closed and DNA topologically bound (Chan *et al.*, 2012; Beckouet *et al.*, 2016). Thus, the Smc3p Mcd1p interface has been postulated to serve as a regulated DNA exit gate and as well as a regulated entrance gate.

While compelling in its simplicity, viewing the function of the Smc3p Mcd1p interface and cohesin regulators solely through the lens of a putative DNA gate has been challenged by several observations. First, cohesin is stably bound to DNA when Smc3p K112 K113 acetylation is prevented by mutating the critical lysines (*smc3 K112R K113R)* (Unal *et al.*, 2008; Rowland *et al.*, 2009). Thus, acetylation of these residues is not needed to stabilize topological entrapment of DNA in vivo. However, both *smc3 K112R K113R* and *eco1*Δ *wpl1*Δ mutants exhibit a dramatic defect in sister chromatid cohesion (Rowland *et al.*, 2009; Sutani *et al.*, 2009; Chan *et al.*, 2012; Guacci and Koshland, 2012). These results suggest that acetylation promotes cohesion by an additional step beyond preventing DNA release through an exit gate. Finally, recent experiments suggest that the function of cohesin and other Smc complexes requires a conformational change of the coiled coil, likely driven by the head ATPase activity (Soh *et al.*, 2015). The binding of Mcd1p to both the head domain of Smc1 and the coiled coil of Smc3 provides a potential mechanism to transduce the ATP state of the head domain to initiate a conformation change of the coiled coils.

Here, we use a fusion protein of Smc3p and Mcd1p to permanently shut the putative DNA gate as a tool to evaluate gate-independent functions of this interface. We examined whether the fusion could suppress the need for the cohesin loader Scc2p, the Scc3p cohesin subunit, Pds5p and Smc3p K112 K113 acetylation. Our results show that in vivo all the regulators of cohesin promote DNA binding or cohesion by mechanisms independent of opening the Smc3p Mcd1p interface. Furthermore, mutations altering the interface reveal it has a gate-independent activity essential for cohesin to bind DNA *in vivo*. These observations suggest new models for the Smc3p Mcd1p interface in topological entrapment and as a target of cohesin regulators.

## Results

We wanted to assess whether cohesin regulators control cohesin function by modulating the transient opening of the Smc3p Mcd1p interface. To do so, we used a previously characterized gene fusion in which the open reading frame of *MCD1* was placed in frame at the end of the open reading frame for *SMC3* (Gruber *et al.*, 2006). The product of this gene fuses the Mcd1p N-terminus to the Smc3p C-terminus so this putative DNA gate cannot open. It supports viability as sole source of both Smc3p and Mcd1p. Consequently, regulators that act solely by stabilizing the exit gate should no longer be needed for cohesin function in fusion bearing strains.

We used the Smc3p Mcd1p fusion to test gate-dependent functions of three proteins associated with the trimer cohesin, the Scc3p cohesin subunit as well as the Scc2p and Pds5p regulators. Structural or functional features of Scc2p, Scc3p and Pds5p suggested that they may act directly or indirectly to modulate the putative Smc3p Mcd1p gate to allow DNA entry or exit. Evidence suggests Scc2p promotes cohesin loading via hinge-hinge dimer opening, so the fusion shouldn’t suppress the essential requirement for Scc2p (Gruber *et al.*, 2006). Scc3p is required for cohesin binding to DNA, cohesion and condensation (Tóth *et al.*, 1999; Roig *et al.*, 2014; Orgil *et al.*, 2015). Scc3p binds to Pds5p and Wpl1p as well as Mcd1p (Rowland *et al.*, 2009; Orgil *et al.*, 2015). Thus, both Scc3p and Pds5p are in a position to modulate the Smc3 Mcd1p interface. If they serve only to keep the interface closed, the fusion should suppress the effects of depleting Scc3p and Pds5p.

To test these predictions, we first built haploid strains bearing the fusion gene as the sole source of both Smc3p and Mcd1p function. We then replaced the endogenous *SCC2, SCC3, PDS5* genes with conditional alleles containing a C-terminal 3V5 tag and an auxin-inducible degron (AID) called *SCC3-AID, SCC2-AID* or *PDS5-AID*. The AID degron enabled the rapid degradation of each of these proteins upon auxin addition (Eng *et al.*, 2014). As a control, we placed the same *AID* constructs in an otherwise wild-type haploid strain to compare the effects of their depletion with and without the fusion.

### Fusion strains still require Scc2p, Scc3p and Pds5p for viability and sister chromatid cohesion

We first assessed whether the fusion bypasses the need for Scc2p, Scc3p and Pds5p for cell viability. Strains with and without the fusion as sole source were grown to saturation and then plated in 10-fold serial dilution either in the presence or absence of auxin. For clarity, strains without fused cohesin will be referred to as normal strains or as having normal cohesin. As expected, depletion of Scc2p-AID, Scc3p-AID or Pds5p-AID were lethal in normal strains (Figures 2A & 2B top panel) (Eng *et al.*, 2014). Depletion of these same proteins in fusion strains was also lethal (Figure 2A & 2B bottom panels). The inviability of fusion strains indicates that Scc2p, Scc3p and Pds5p are required for an essential cohesin biochemical activity other than blocking exit through the putative Smc3p Mcd1p gate.

**Figure 2:**
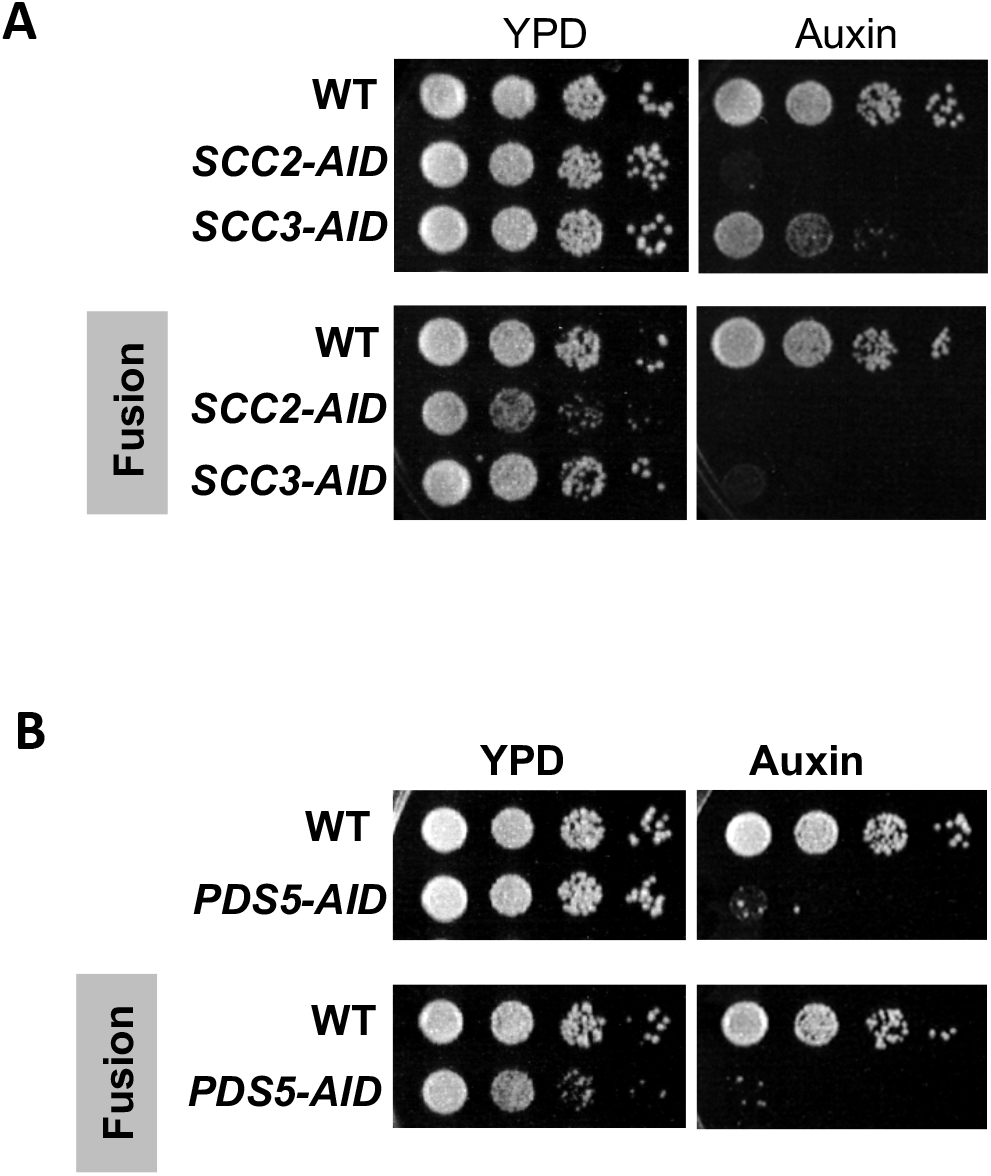
Cohesin regulators are required fusion cohesin function. (A-B) Haploid strains with normal cohesin or fusion were grown to saturation, then plated at 10-fold serial dilution onto YPD alone or containing auxin (750μM) and incubated at 23°C for 3d. (A) Scc2p-AID and Scc3p-AID depletion causes in inviability in normal and fusion cohesion strains. Top panel shows strains with normal cohesin, WT (VG3620-4C), containing *SCC2-3V5-AID* (VG3630-7A) or *SCC3-3V5-AID* (VG3808-1A). Bottom panel shows strains with fusion cohesin, WT (VG3940-2D) or with *SCC2-3V5-AID* (VG3945-1A) or *SCC3-3V5-AID* (VG3946-7B). (B) Pds5p-AID depletion causes in inviability in normal and fusion cohesion strains. Top panel shows strains with normal cohesin, WT (VG3620-4C) or containing *PDS5-3V5-AID2* (VG3954-10C). Bottom panel shows strains with fusion cohesin, WT (VG3940-2D) or containing *PDS5-3V5-AID2* (VG3955-4D).

To begin to understand the exit gate-independent molecular functions of Scc2p, Scc3p and Pds5p, we assayed sister chromatid cohesion in our AID strains with and without the fusion. We used a regimen of synchronous arrest in mid-M phase as follows. Cultures were arrested in G1 where the AID tagged proteins depleted. Cells were released from G1 into media containing auxin and nocodazole to re-arrest cells in mid-M phase under AID depletion conditions (Materials and methods & Suppl Figure 1A). Cell cycle arrest and depletion of AID tagged proteins were confirmed by FACS (Suppl Figures 1B and 2A) and by Western (Suppl Figures 1C, 1D and 2B). Mid-M arrested cells were assayed for cohesion at a *CEN*-distal (*LYS4*) locus on chromosome IV using the GFP-LacI and LacO system (Materials and methods). In budding yeast, after replication the sister chromatids are so closely associated they cannot be resolved as individual chromatids by fluorescence microscopy (Guacci *et al.*, 1994). Therefore, in the LacO-LacI assay, haploids in mid-M exhibit a single GFP focus whereas cells that are defective for cohesion have two GFP spots, one from each sister chromatid.

Mid-M arrested cells with normal cohesin have cohesion showed two GFP spots in only 5-10% of cells, whereas cells depleted of Scc2p-AID, Scc3p-AID, or Pds5p-AID had a major cohesion defect as evidenced by more than 75% two GFP spots, consistent with previous results (Figures 3A & 3B, top panels) (Eng *et al.*, 2014). Most mid-M arrested cells with fusion cohesin had cohesion although a modest cohesion defect was observed (∼25% cells with two GFP spots) similar to that seen in previous studies (Figures 3A & 3B, bottom panels) (Gruber *et al.*, 2006; Bloom *et al.*, 2018). In contrast, fusion cells depleted for Scc2p, Scc3p or Pds5p had severe cohesion defects similar in magnitude to that seen after depletion in cells with normal cohesin (Figures 3A & 3B, bottom panels). These results show that the all three proteins play critical roles in sister chromatid cohesion independent of any putative role in keeping the Smc3p Mcd1p interface closed.

**Figure 3:**
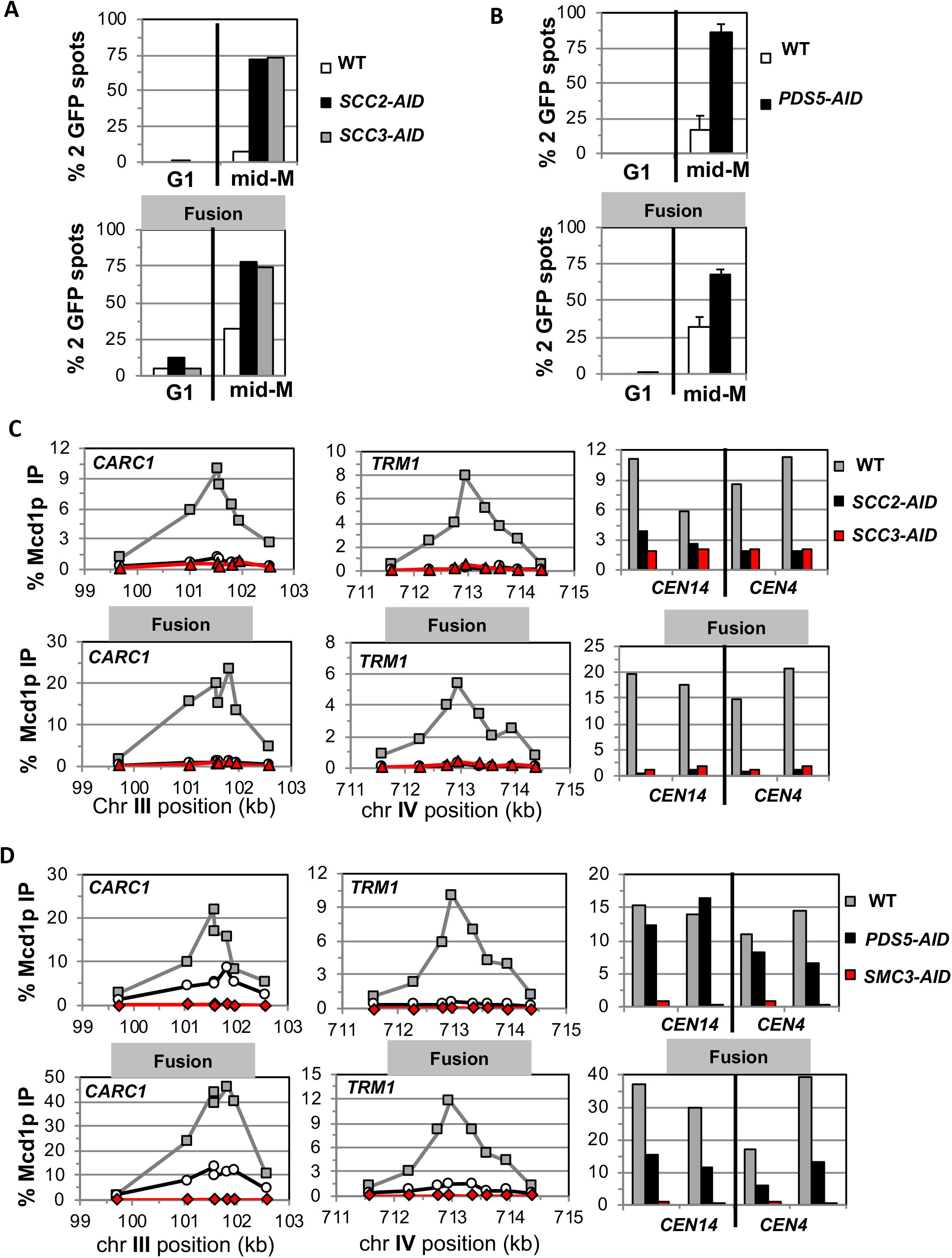
Fusion cohesin requires cohesin regulators for sister chromatid cohesion and cohesin binding to DNA. Haploids strains with normal cohesin or fusion cohesin from figure 2 were grown to mid-log phase then arrested in G1 using αfactor, auxin added to induce loss of AID tagged proteins then released into media containing nocodazole and auxin to arrest in mid-M phase under AID depletion conditions as described for synchronous mid-M phase arrest in materials and methods. Cells were fixed and processed to monitor loss of sister chromatid cohesion and for ChIP to monitor cohesin DNA binding. (A-B) Cohesion loss monitored in mid-M phase cells. The number of GFP spots was scored in G1 arrested cells and mid-M phase cells. The percentage of cells with 2 GFP spots was plotted. 100-200 cells were scored for each data point and data was generated from 2 independent experiments. (A) Scc2p-AID or Scc3p-AID depletion induces cohesion loss in strains with normal or fusion cohesin. Top panel is strains with normal cohesin and bottom panel is fusion cohesin strains. WT (White), *SCC2-AID* (black) and *SCC3-AID* (grey). (B) Pds5p-AID depletion induces cohesion loss in strains with normal or fusion cohesin. Top panel is strains with normal cohesin and bottom panel is fusion cohesin strains. WT (White) and *PDS5-AID* (black). (C-D) mid-M phase arrested cells fixed and processed for ChIP using anti-Mcd1p antibodies as described in materials and methods. Mcd1p binding was assessed by qPCR and presented as percentage of total DNA using the same primer pairs at each site. Left Panel is chromosome III peri-centric region (*CARC1*), middle panel is chromosome IV arm CAR region (*TRM1*) and right panel is regions immediately adjacent to *CEN4* and *CEN14*. (C) Scc2p-AID or Scc3p-AID depletion induces loss of fusion cohesin and normal cohesin binding to DNA. Top panel is strains with normal cohesin and bottom panel is fusion cohesin strains. WT (grey), *SCC2-AID* (black) and *SCC3-AID* (red). (D) Pds5p-AID depletion reduces the amount of fusion cohesin and normal cohesin binding to DNA. Top panel is strains with normal cohesin and bottom panel is fusion cohesin strains. WT (grey), *PDS5-AID* (black) and *SMC3-AID* (red).

### The Smc3 Mcd1p fusion does not suppress the effects of Scc2p, Scc3p and Pds5p depletion on cohesin binding to DNA

To understand further the molecular basis for the gate-independent functions of cohesin regulators, we compared the effect of Scc2p-AID and Scc3p-AID depletion on normal and fusion cohesin binding to chromosomes in cells subjected to our regimen of synchronous arrest. These cells were subjected to chromatin immunoprecipitation (ChIP) using anti-Mcd1p antibodies (Materials and methods). In wild-type cells, fusion cohesin bound with the same pattern as normal cohesin at a pericentric *CAR* (*CARC1*) and an arm *CAR* (*TRM1*) locus as well as immediately adjacent to *CEN4* and *CEN14*. (Figure 3C). Thus, the minor defect in cohesion observed in the fusion strain was not due to effects on fusion cohesin levels or distribution on chromosomes. For both normal and fusion cohesin, depletion of Scc2p-AID or Scc3p-AID completely abolished cohesin binding at peri-centric *CARC1* and arm CAR *TRM1* and dramatically reduced binding near *CEN4* and *CEN14* (Figure 3C). The lack of cohesin binding to DNA was not due to loss of either Mcd1p or Smc3p Mcd1p fusion as shown by Western Blot analysis (Suppl Figure 2E). Thus, Scc2p and Scc3p are essential for both normal cohesin and fusion cohesin to bind chromosomes. The robust fusion cohesin binding to DNA means the Smc3p Mcd1p interface is not an entry gate for DNA. Therefore, Scc2p and Scc3p must promote cohesin DNA binding through an activity independent of preventing DNA entry through the putative Smc3p Mcd1p gate.

We also assessed the effect of Pds5p-AID depletion on cohesin binding in mid-M arrested cells bearing normal cohesin or with fusion cohesin. Pds5p-AID depletion was confirmed by both Western Blot and loss of Pds5p binding to chromosomes as assayed by ChIP (Suppl Figures 2B & 2D). There was a decrease in Mcd1p of normal cohesin as well as fusion protein, albeit more modestly than Mcd1p (Suppl Fig 2C). This result is consistent with Pds5p serving partially to protecting Mcd1p from factors that degrade it (Stead *et al.*, 2003; D’Ambrosio and Lavoie, 2014). At the pericentric *CARC1*, Pds5p-AID depletion reduced normal and fusion cohesin by 60%-75%, but the reduction for both was more severe at *TRM1* (Figure 3D). Adjacent to *CENs*, normal cohesin binding was unperturbed whereas fusion cohesin binding was reduced ∼50%. Thus, the fusion did not suppress the decrease cohesin binding to DNA caused by Pds5p-AID depletion. These results show that Pds5p promotes cohesion and cohesin binding to DNA by a mechanism other than preventing DNA exit through the putative Smc3p Mcd1p gate.

### Smc3 head acetylation in the fusion is required for efficient generation of sister chromatid cohesion but not for cell viability

The exit gate model posited that Wpl1p opens the Smc3p Mcd1p interface to allow DNA escape from cohesin, but gate opening is blocked by Eco1p mediated acetylation of Smc3p at K112 K113 (Chan *et al.*, 2012; Beckouet *et al.*, 2016). Loss of Eco1p function is lethal and causes defects in cohesion and condensation in budding yeast (Skibbens *et al.*, 1999; Tóth *et al.*, 1999; Bloom *et al.*, 2018). Previous work showed that the fusion suppressed the lethality and condensation defects of *eco1*Δ (Chan *et al.*, 2012; Bloom *et al.*, 2018). However, the *eco1*Δ fusion strain had an increased defect in sister chromatid cohesion compared to the wild-type fusion (Bloom *et al.*, 2018). Eco1p acetylates Smc3p at multiple sites (Unal *et al.*, 2008). We wanted to see whether the partial suppressive effects of the fusion were due solely to acetylation at the essential K112 K113 residues.

We addressed this question by placing the *smc3 K112R K113R* (RR) mutant in the fusion and assessing the effect on cohesin function. If Smc3p K112 K113 acetylation function solely to keep this interface closed, then it should be completely suppressed by the fusion. Strains bearing the RR fusion as the sole Smc3p and Mcd1p source is viable but is cold sensitive and sensitive to the microtubule destabilizing drug benomyl (Figures 4A & 4B). Thus, the fusion suppressed the essential function of Smc3p head acetylation, just as it suppressed an *eco1*Δ, but defects in cohesin function remained.

**Figure 4:**
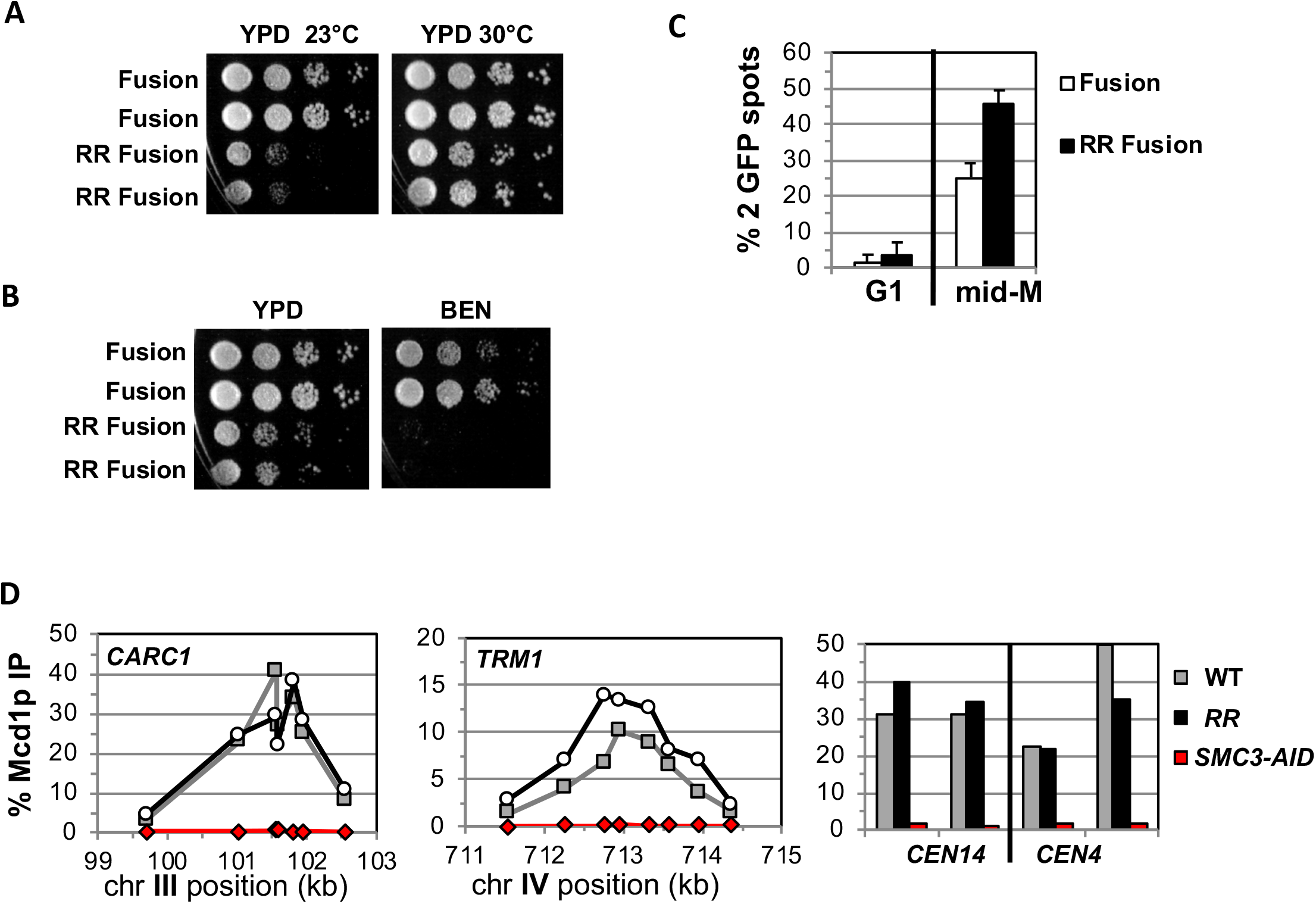
Fusion cohesin with the smc3 K112R K113R mutations are viable but have defects in growth defects in both growth and sister chromatid cohesion. (A-B) Fusion cohesion bearing *smc3-K112R, K113R* (RR) mutations are viable but cold sensitive and benomyl sensitive. Haploid strains with wild-type fusion cohesin (VG3940-2D) or RR fusion (VG3930-5C) were grown to saturation at 30°C then plated at 10-fold serial dilution onto (A) YPD at either 23°C or 30°C and incubated for 3d. (B) YPD alone or YPD containing benomyl at 12.5ug/ml (BEN) then incubated for 23°C for 4d. (C-D) Strains in A-B were synchronously arrested in mid-M as described in figure 3 above except cells were grown at 30°C and auxin was omitted. Cells were fixed and processed to assess cohesion and for ChIP. (C) The RR mutation in fusion cohesin causes an increased defect in cohesion fusion. Cohesin was scored and plotted as described in figure 3. Fusion cohesin (White) and RR fusion cohesin (black). (D) Fusion and RR fusion cohesin bind DNA at similar levels. mid-M phase arrested cells were fixed and processed for ChIP using anti-Mcd1p antibodies as described in materials and methods and figure 3. Left Panel is chromosome III peri-centric region (*CARC1*), middle panel is chromosome IV arm CAR region (*TRM1*) and right panel is regions immediately adjacent to *CEN4* and *CEN14*. WT fusion (grey), RR fusion (black) and *SMC3-AID* (red).

We also examined sister chromatid cohesion and cohesin binding to chromosomes in fusion wild-type and RR strains. Cells were synchronously arrested in mid-M phase as described for AID depletion experiments above except auxin was omitted and cells grown at 30°C due to the severe growth defect of RR fusion strains at 23°C. Forty-five percent of the *s*mc3p-RR fusion bearing cells had a cohesion defect, an ∼2 fold increase compared to wild-type fusion cells (Figure 4C). This increase suggests that Smc3 head acetylation regulates cohesion by a mechanism other than blocking DNA exit through the putative gate. The cohesion defect could also explain the benomyl sensitivity as microtubule destabilization would abrogate early S phase bipolar attachments necessary for proper segregation in cells with severe cohesion defects (Guacci and Koshland, 2012). Finally, wild-type fusion and smc3p-RR fusion cohesin bound to chromosomes at similar levels and these fusion proteins are also present in cells at similar levels (Figure 4D, Suppl Fig 3B). This similarity means the increased cohesion defects in the RR fusion was not due to decreased cohesin binding to DNA. These results indicate that Smc3p head acetylation promotes efficient sister chromatid cohesin independent of a putative function in keeping Smc3p Mcd1p interface closed.

### Smc3p Mcd1p interface controls the integrity of another cohesin interface to allow cohesin binding to DNA

Having established that key cohesin regulators function independently of preventing DNA exit through the Smc3p Mcd1p interface, we turned to characterize the function of the interface itself. Mutations in Mcd1p or Smc3p that affect residues in the interface are lethal (Arumugam *et al.*, 2006; Eng *et al.*, 2014; Gligoris *et al.*, 2014; Robison *et al.*, 2018). One such mutation, *smc3-L1029R* was thought to prevent cohesin binding to DNA because when tagged with GFP, it failed to immunolocalize to the cohesin-rich centromere cluster (Gligoris *et al.*, 2014). This mutation also abolished a chemical crosslink normally detected between Smc3p and Mcd1p (Gligoris *et al.*, 2014). These results were interpreted to mean that interface mutants eliminated Smc3p Mcd1p association, thereby allowing DNA to escape from cohesin entrapment. However, our results with fusion cohesin show that Pds5p and Scc3p promote DNA binding by mechanisms independent keeping this interface “gate” closed. Their binding proximal to the interface raised the alternative possibility that the interface also promoted a gate-independent biochemical property to regulate cohesin binding to DNA.

To test this possibility, we examined mutations that alter amino acids of the interface from both Mcd1p (*mcd1-L75K* and *mcd1-L89K*) and Smc3p (*smc3-I1026R* and *smc3-L1029R*). All four mutations were previously shown to be unable to support viability (Arumugam *et al.*, 2006; Gligoris *et al.*, 2014). Three possible outcomes are expected from the fusion mutants. If all these mutations abrogate cohesin function solely by preventing closure of a putative Smc3p Mcd1p interface exit gate, then all four mutants should be suppressed by the fusion. Alternatively, if the fusion fails to suppress all these mutants, then these residues must modulate cohesin in a way distinct from keeping a putative exit gate closed. Finally, the fusion could suppress mutants in only Smc3p or only Mcd1p. For example, if Mcd1p residues only promote association with Smc3p whereas Smc3p residues have a second function, then the fusion would only suppress only the *mcd1* mutations.

Before testing these mutants in the fusion, we further characterized them in normal cohesin to provide a baseline for judging their effects on fusion cohesin. The *smc3* mutants were placed into a haploid strain bearing *SMC3-3V5-AID* (*SMC3-AID*) as the sole *SMC3* whereas the *mcd1* mutants were placed in a haploid bearing *MCD1-AID* as the sole *MCD1*. The corresponding wild-type alleles were introduced into the *AID* strains as positive controls and the AID alone served as the negative control. Strains were tested by dilution plating on auxin containing media to assess viability under *AID* depletion conditions. As expected, *SMC3-AID* alone or *MCD1-AID* alone were inviable on auxin whereas those also containing wild-type alleles are viable (Figures 5A & 5B and (Eng *et al.*, 2014). None of the mutant alleles rescued the inviability of the *AID* strains, corroborating previous studies that these interface residues provide an essential cohesin function (Figures 5A & 5B).

**Figure 5:**
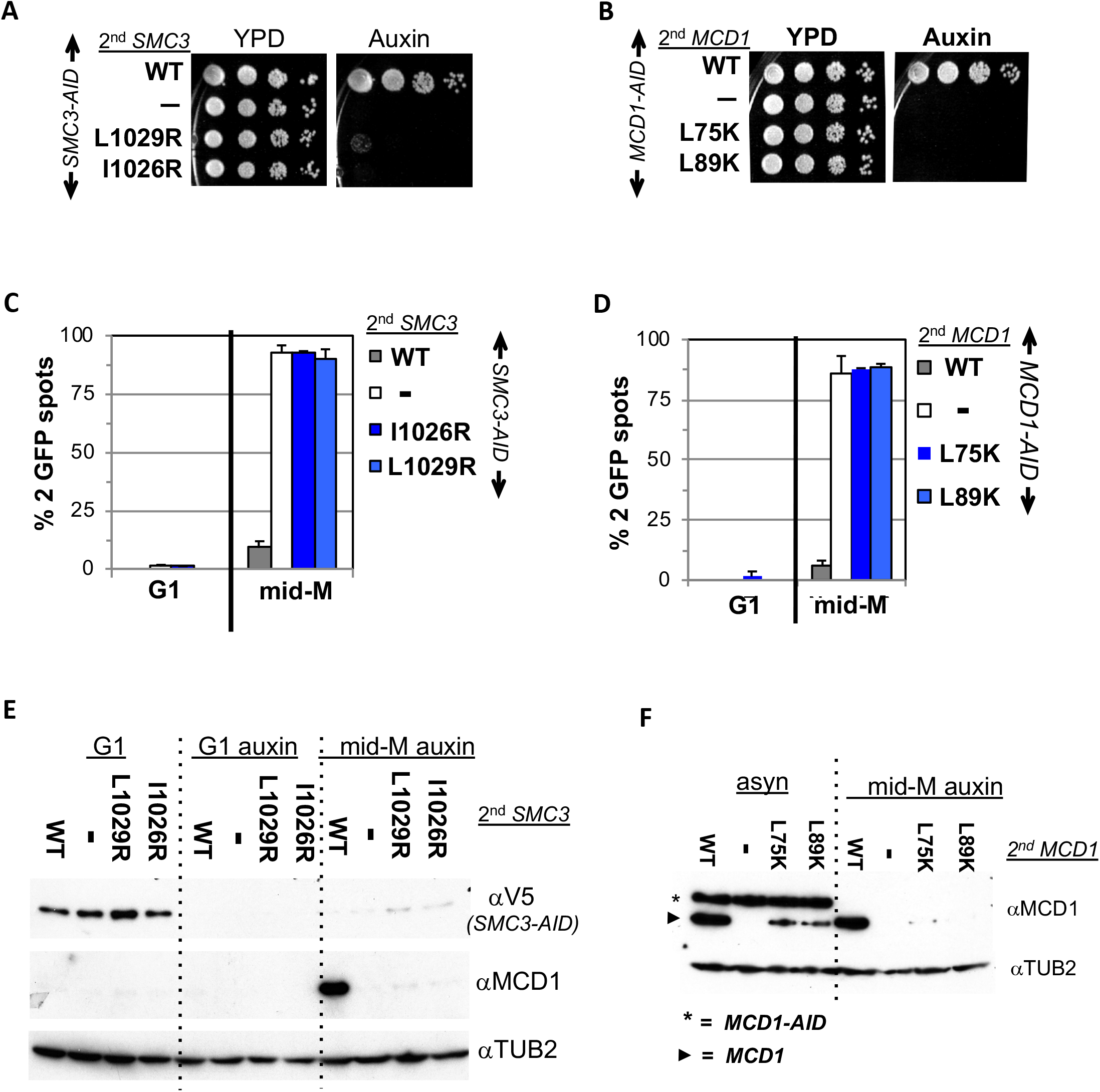
Smc3p coiled-coil and Mcd1p NHD interface residues are required for cohesin function and integrity. (A) Viability loss of Smc3p interface mutant strains. Haploid *SMC3-AID* strain alone (VG3651-3D) or containing either wild-type (WT; BRY474), *smc3-I1026R* (VG3905-7A) or *smc3-L1029R* (BRY492) were grown and plated as described in Figure 2. (B) Viability loss of Mcd1p interface mutant strains. Haploid *MCD1-AID* strain alone (VG3902-3A) or containing, either wild-type (WT; VG3914-2C), *mcd1-L75K* (VG3916-5B) or *mcd1-L89K* (VG3918-9D) were grown and plated as described in Figure 2. (C-D) Haploids in A & B were synchronously arrested in mid-M phase as described in Figure 3. The number of GFP spots was scored in G1 arrested cells and mid-M phase cells. The percentage of cells with 2 GFP spots was plotted. 100-200 cells were scored for each data point and data was generated from 2 independent experiments. (C) Cohesion loss in *smc3* interface mutant mid-M cells. *SMC3-AID* alone (white), or containing WT (grey), *smc3-I1026R* (dark blue) and *smc3-L1029R* (light blue). (D) Cohesion loss in *mcd1* mutant mid-M cells. *MCD1-AID* strain alone (white) or containing, either WT (grey), *mcd1-L75K* (dark blue) or *mcd1-L89K* (light blue). (E-F) Protein extracts from the synchronous arrest regimen in A & B were made from G1 cells, auxin treated G1 cells and mid-M cells then subjected to Western blot analysis. (E) Mcd1p is degraded in mid-M phase *smc3* interface mutants. Top panel detects Smc3p-3V5-AID (αV5), middle panel detects Mcd1p (αMcd1). Tubulin (αTub2p; bottom panel) was used as a loading control. As a control we showed in Supplemental Figure 4E that the *smc3-L1029R* mutation did not affect its protein levels by comparing 6HA epitope tagged Smc3p (VG3943-1C) and Smc3p-L1029R (VG3944-3D). (F) Mcd1p interface mutants are degraded in mid-M cells. Top panel detects Mcd1p (αMcd1) with star indicating Mcd1p-AID and arrow head Mcd1p WT and interface mutants, bottom panel detects Tubulin (αTub2p).

We next examined whether the four interface mutants compromised sister chromatid cohesion. Cells were synchronously arrested in mid-M phase under AID depletion conditions and arrest confirmed by FACS (Materials and Methods; Suppl Figures 4A & 4B). We assessed cohesion at the *LYS4* locus using the LacO-GFP, LacI system. As expected, almost all *SMC3-AID* and *MCD1-AID* cells had 2 GFP spots indicating a total loss of cohesion whereas cells with the respective wild-type alleles promote cohesion so few cells had 2 GFP (Figures 5C & 5D). All four mutants alleles were totally defective in cohesion, similar to what was seen in the *AID* alone strains (Figures 5C & 5D).

To investigate why this cohesin defect was so severe, we examined whether these mutants support the integrity of the cohesin trimer. Mcd1p is not detectable from anaphase through G1 due to degradation but is present from the S phase through mid-M. However, Mcd1p presence from S phase through mid M is dependent upon its binding to both Smc1p and Smc3p as Mcd1p is rapidly degraded if either Smc1p-AID or Smc3p-AID are depleted (Çamdere *et al.*, 2015; Guacci *et al.*, 2015). Therefore, Mcd1p presence serves as a readout for the in vivo presence of cohesin trimer (Smc3p, Mcd1p, Smc1p).

As expected, Mcd1p was absent in all G1 cells and Smc3p-AID depleted in all auxin treated cells (Figure 5E). In mid-M cells, Mcd1p was observed when Smc3p (*SMC3 SMC3-AID*) but absent in the *SMC3-AID* alone strain. Mcd1p was also absent in smc3p-AID mid-M cells expressing smc3p-L1029R or smc3p-I1026R (Figure 5E). Similarly, Mcd1p-AID was depleted in all mid-M phase cells and wild-type Mcd1p is present whereas the mcd1p-L75K and mcd1p-L89K mutants were absent (Figure 5F, right side). Our results show that perturbation of the Smc3p Mcd1p interface with mutations in either Smc3p or Mcd1p led to Mcd1p degradation. Interestingly, the degradation of the Mcd1p NHD mutants was partially suppressed by the presence of the Mcd1p-AID, providing an explanation why this degradation was missed previously. This suppression can be explained by interallelic complementation (Eng *et al.*, 2015)

The Mcd1p degradation explained the loss of all cohesin function in the *smc3* and *mcd1* mutants but also complicated interpretations of the function of the interface. These mutations could indeed abrogate stable association of Smc3p and Mcd1p leading to Mcd1p degradation, or simply alter the interface in a way that allowed protease accessibility to the Mcd1p N terminus leading its degradation. In either case, the loss of Mcd1p in the normal strains precluded determining whether the interface performs another function distinct from an exit gate.

With this foundation we examined the properties of fusions containing either *smc3*-*I1026R* or *smc3*-*L1029R* in an *SMC3-AID* background, and either *mcd1-L75K* or *mcd1-L89K* mutations in an *MCD1-AID* background. These strains allowed us to examine the properties of the mutated fusions in the absence of the corresponding wild-type normal (unfused) subunit after auxin addition. Dilution plating onto auxin containing media revealed that the fusion failed to suppress the inviability of any of the *smc3* or *mcd1* interface mutants (Figures 6A and 7A). We then tested whether the fusion suppressed the *smc3* and *mcd1* interface mutant induced degradation of Mcd1p. All four mutant fusion proteins were present in mid-M cells, although the mcd1p fusion mutants were slightly less abundant than the wild type fusion (Figures 6B & 7B). The presence of the mutant fusion proteins demonstrated that the complete Mcd1p loss observed when normal cohesin contains interface mutations depended upon a free N terminus of Mcd1p. More importantly, the inviability of the fusion mutants was not due to absence of fusion protein, but rather due to some other biochemical defect caused by the mutations.

**Figure 6:**
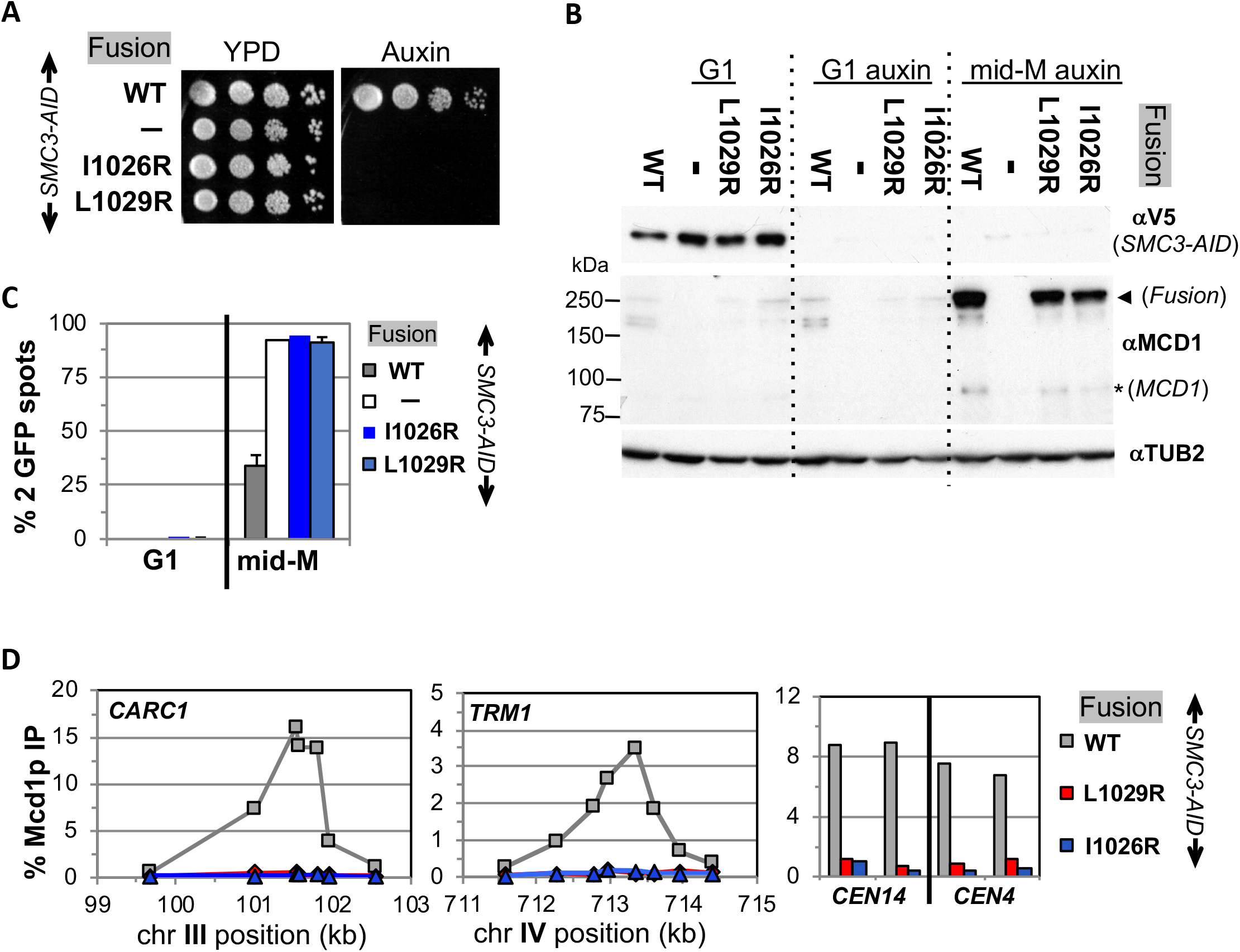
Fusion cohesin requires Smc3p coiled coil residues for function. (A) Viability loss of Smc3p coiled coil mutant fusion strains. Haploid *SMC3-AID* strain alone (VG3651-3D) or containing fusion cohesin, either wild-type (WT; VG3694-7C), *smc3-I1026R* (VG3908-17B) or *smc3-L1029R* (VG3872-3B) were grown and plated as described in Figure 2. (B-D) Haploids in A were synchronously arrested in mid-M phase as described in Figure 3. (B) Fusion WT and *smc3* interface mutant proteins are present in mid-M cells. Protein extracts from G1 and auxin treated G1 and mid-M phase arrested cells were subjected to Western blot analysis using antibodies against V5 to detect Smc3p-3V5-AID (αV5; top panel) and against Mcd1p (αMcd1p; middle panel). Arrow head indicates fusion protein and star (*) indicates normal Mcd1p. Tubulin (αTub2p; bottom panel) was used as a loading control. (C) Cohesion is completely lost in *smc3* mutant fusion mid-M phase arrested cells. The number of GFP spots was scored in G1 arrested cells and mid-M phase cells. The percentage of cells with 2 GFP spots was plotted. *SMC3-AID* alone (white), or containing fusion WT (grey), *smc3-I1026R* (dark blue) and *smc3-L1029R* (light blue). 100-200 cells were scored for each data point and data was generated from 2 independent experiments. (D) fusion *smc3* interface mutant cohesin fails to bind DNA in mid-M phase. Fusion cohesin binding to chromosomes was assayed by ChIP using anti-Mcd1p antibodies as described in figure 3. WT fusion (grey), *smc3-I1026R* fusion (blue) and *smc3-L1029R* fusion (red).

**Figure 7:**
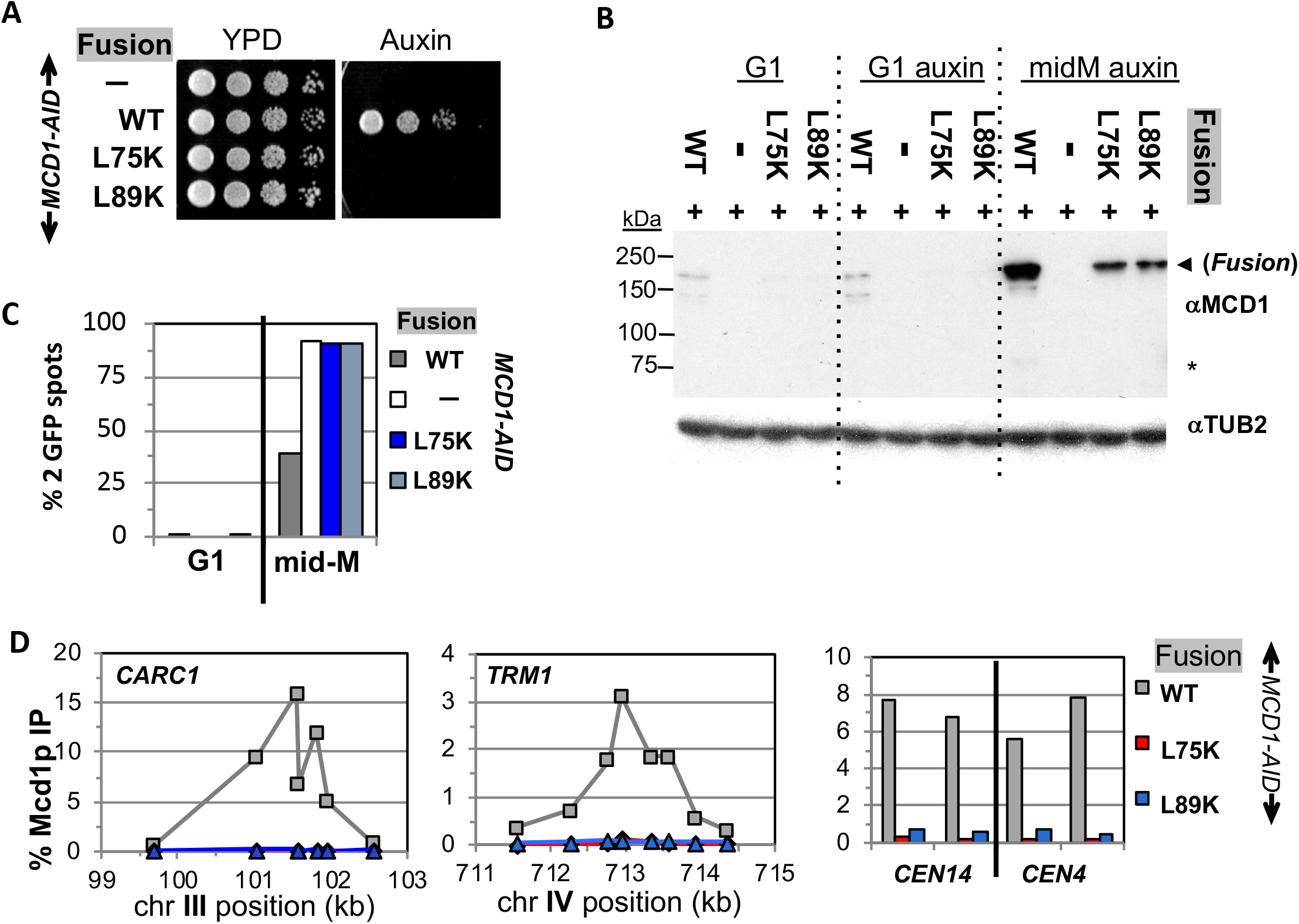
Fusion cohesin requires Mcd1p NHD residues for function. (A) Viability loss of Mcd1p NHD mutant fusion strains. Haploid *MCD1-AID* strain alone (VG3902-3A) or containing fusion cohesin, either wild-type (WT; VG3937-2C), *mcd1-L75K* (VG3938-3A) or *mcd1-L89K* (VG3939-7B) were grown and plated as described in Figure 2. (B-D) Haploids in (A) were synchronously arrested in mid-M phase as described in Figure 3. (B) Fusion WT and mcd1 NHD mutant proteins are present in mid-M cells. Protein extracts from G1, auxin treated G1 and mid-M phase cells were subjected to Western blot analysis using antibodies against Mcd1p (αMcd1p; top panel). Arrow head indicates fusion protein and star (*) indicates normal Mcd1p. Tubulin (αTub2p; bottom panel) was used as a loading control. (C) Cohesion is completely lost in *mcd1* mutant fusion mid-M cells. The number of GFP spots was scored in G1 arrested cells and mid-M phase cells. The percentage of cells with 2 GFP spots was plotted. Haploid *MCD1-AID* strain alone (white) or containing fusion cohesin, either WT (grey), *mcd1-L75K* (dark blue) or *mcd1-L89K* (light blue). 100-200 cells were scored for each data point. (D) fusion *mcd1 NHD* mutant cohesin fails to bind DNA in mid-M phase. Fusion cohesin binding to chromosomes was assayed by ChIP using anti-Mcd1p antibodies as described in figure 3. WT (grey), *mcd1-L75K* (red) and *mcd1-L89K* (blue).

We then examined the four mutant fusions for sister chromatid cohesion and cohesin binding to chromosomes in cells synchronously arrested in mid-M under auxin depletion conditions. All four mutant fusion strains had almost no sister chromatid cohesion when depleted for Mcd1p-AID or Smc3p-AID (Figures 6C and 7C). By ChIP, all four fusion mutants exhibited complete loss of cohesin binding to *CARs* and *CEN* DNA when depleted for Mcd1p-AID or Smc3p-AID (Figures 6D and 7D). The loss of cohesin binding to DNA explains why the interface fusion mutants were inviable and unable to generate cohesion.

The shared dramatic phenotypes for all four mutant fusions lead to several important conclusions about the Smc3p Mcd1p interface. First, the amino acids of Mcd1p and Smc3p that make up the interface must have essential biochemical activities other than directly maintaining the topological integrity of the cohesin ring at this interface. Second, the fact that mutations in Smc3p Mcd1p interface residues have the same phenotypes suggests that these Smc3p and Mcd1p regions collaborate to perform the same biological function, likely through their interaction. Finally, the absence of DNA binding in the mutant fusions suggests that at least one essential biochemical activity of the interface is to promote DNA entrapment by modulating the opening or stable closing of a distal interface of cohesin. This modulation of a distal interface could be achieved directly through changing cohesin structure or indirectly through a regulator.

## Discussion

Elucidating how cohesin carries out its many biological activities requires understanding how cohesin’s topological entrapment of DNA is established, maintained and released. A reversible dissociation of the interface between Smc3p and Mcd1p has been proposed to be either an entrance gate or an exit gate enabling cohesin to establish or maintain topological DNA entrapment, respectively (Chan *et al.*, 2012; Murayama and Uhlmann, 2015; Beckouet *et al.*, 2016). Here we assessed gate-independent functions of this interface in yeast using a fusion protein where Smc3p at its C-terminus was fused to Mcd1p at its N-terminus. Our results reveal surprising phenotypes that provide significant new insights into the mechanism and regulation of topological entrapment.

The fact that fusion cohesin and normal unfused cohesin bind DNA with the same pattern and at similar levels demonstrates that the Smc3p Mcd1p interface is not a direct entrance gate (this study). However, we show that this interface has a gate-independent function for DNA binding since the interface mutants in the fusion fail to bind DNA. Studies of fluorescent labeled cohesin in vivo reveals two pools of cohesin that bind DNA either unstably or stably (Gerlich *et al.*, 2006; Chan *et al.*, 2012). Both states are captured by ChIP as evidenced by the fact that *eco1* mutants dramatically reduces the amount of cohesin in the stable state without altering the amount bound to chromosomes (Lengronne *et al.*, 2006; Noble *et al.*, 2006). The failure to detect chromosome binding of cohesins with the fusion interface mutations by ChIP suggests that these mutant cohesins cannot even establish unstable topological entrapment. The simplest explanation for the very stringent function for the interface in DNA binding is that it must promote the opening or closing of a distal interface of the cohesin trimer to initiate stable DNA binding.

Crosslinking studies suggested that the hinge dimer interface opens to enable DNA entry (Gruber *et al.*, 2006). Therefore, our results suggest that the Smc3p Mcd1p interface likely controls the hinge-hinge dimer. The control could be indirect through the recruitment of factors like Scc2p or Scc3p, which are essential for cohesin loading onto DNA. This model would predict that a fusion with the *smc3-L1029R* mutation would fail to coimmunoprecipitate with Scc2p and Scc3p which is not the case (Suppl Figure 5), disfavoring this model. Therefore, we prefer the idea that Scc2p and Scc3p may modulate the interface which in turn opens or closes a distal interface.

How could the Smc3p Mcd1p interface regulate a distal interface like the hinge? This interface could directly induce a conformational change in cohesin to alter the distal interface. The existence of different conformations of cohesin (rings, rods and folded rings) have been demonstrated by microscopic techniques like EM and AFM (Onn *et al.*, 2008). Recent work suggests that the cohesin Smc ATPase activity alters head-head interactions and promotes a conformation change that enables DNA binding and cohesion (Çamdere *et al.*, 2015; Huber *et al.*, 2016; Çamdere *et al.*, 2018). Since Mcd1p binds both to the base of the Smc3p coil and the head, it could transduce ATPase dependent changes in the head to the coil, which could cause interconversion of rings to rods or rings to folded rings, thereby affecting the opening or closing of a distal interface.

Our studies also provide important insights into the previously established roles of Pds5p and Smc3p acetylation in cohesin function after DNA binding. Here, we show that these factors must have functions independent of modulating DNA release through the Smc3p Mcd1p interface since the fusion fails to suppress any phenotypes of Pds5p depletion and only partially suppresses the phenotypes of Smc3p lacking K112, K113 acetylation. The post-DNA binding functions of Pds5p and Smc3p acetylation could be totally independent of the interface.

However, Smc3p ATPase domain mutations that bypass acetylation both stabilize the interface and promote cohesin function after DNA binding (Çamdere *et al.*, 2015; Beckouet *et al.*, 2016; Elbatsh *et al.*, 2016). These results suggest that acetylation promotes post DNA binding functions of cohesin by stabilizing the interface. Interestingly, Pds5p which is needed for post DNA binding functions of cohesin is also required for efficient Smc3 acetylation (Chan *et al.*, 2013). Indeed, the fusion likely acts as a partial surrogate for acetylation (partial suppression of acetylation defects) because it enhances the formation of the interface by virtue of holding the Mcd1 NHD and Smc3 coiled coil in close proximity or constrains its conformation. Finally, the establishment of cohesion is normally restricted to S phase. However, two mutations in the Mcd1p NHD (*mcd1-S83D* and *mcd1-K84Q*) mimic respectively phosphorylation by Chk1 kinase and Eco1 acetylation. These interface mutations induce DNA-bound cohesin in mid-M to become cohesive (Heidinger-Pauli *et al.*, 2009). Taken together, all these results suggest that cohesin regulators modulate the interface to promote post DNA binding steps important for cohesion.

We propose an alternative model to explain how Scc2p, Scc3p, Pds5p and Smc3p acetylation impact cohesin DNA binding and cohesin function. Scc2p stimulates cohesin ATPase activity and loading cohesin onto DNA, and Scc3p is required for both Scc2p mediated effects (Murayama and Uhlmann, 2013). We propose Scc2p, Scc3p and cohesin ATPase act through the Smc3p Mcd1p interface to stimulate opening or closing of distal interface to enable establishment of topological entrapment (Figure 8). If the DNA entry is at the hinge, this would stiffen the coiled coils to convert the open ring into a rod-like conformation. Alternatively, conversion into a folded ring could bring the hinges into close proximity to the cohesin loader bound near the Smc heads where loader directly stimulates hinge opening. Our observations also suggest an alternative model for the maintenance of cohesin DNA binding. Wpl1p also acts at the Smc3p Mcd1p interface to release DNA but does so by destabilizing a distal interface (Figure 8 bottom). Pds5p and Smc3p acetylation counter Wpl1p at the Smc3p Mcd1p interface to stabilize DNA binding and promote post-DNA binding steps necessary for cohesion. Distinguishing between our alternative model and the exit gate model will require additional experiments. However, it is worth noting that Pds5p binds Mcd1p near the NHD and the Smc3p coiled coil immediately adjacent to the Smc3p Mcd1p interface (Chan *et al.*, 2013; Eng *et al.*, 2014; Huis in ‘t Veld *et al.*, 2014). Consequently, Pds5p provides a second link between Mcd1p and Smc3p distinct from the interface. This second link could block DNA escape from the trimer should the association between Mcd1p NHD and the Smc3p coiled coil fail transiently or even completely. As such, DNA release from cohesin may be achieved more simply by opening a distal interface than removing both of the links between Mcd1p and Smc3p.

**Figure 8:**
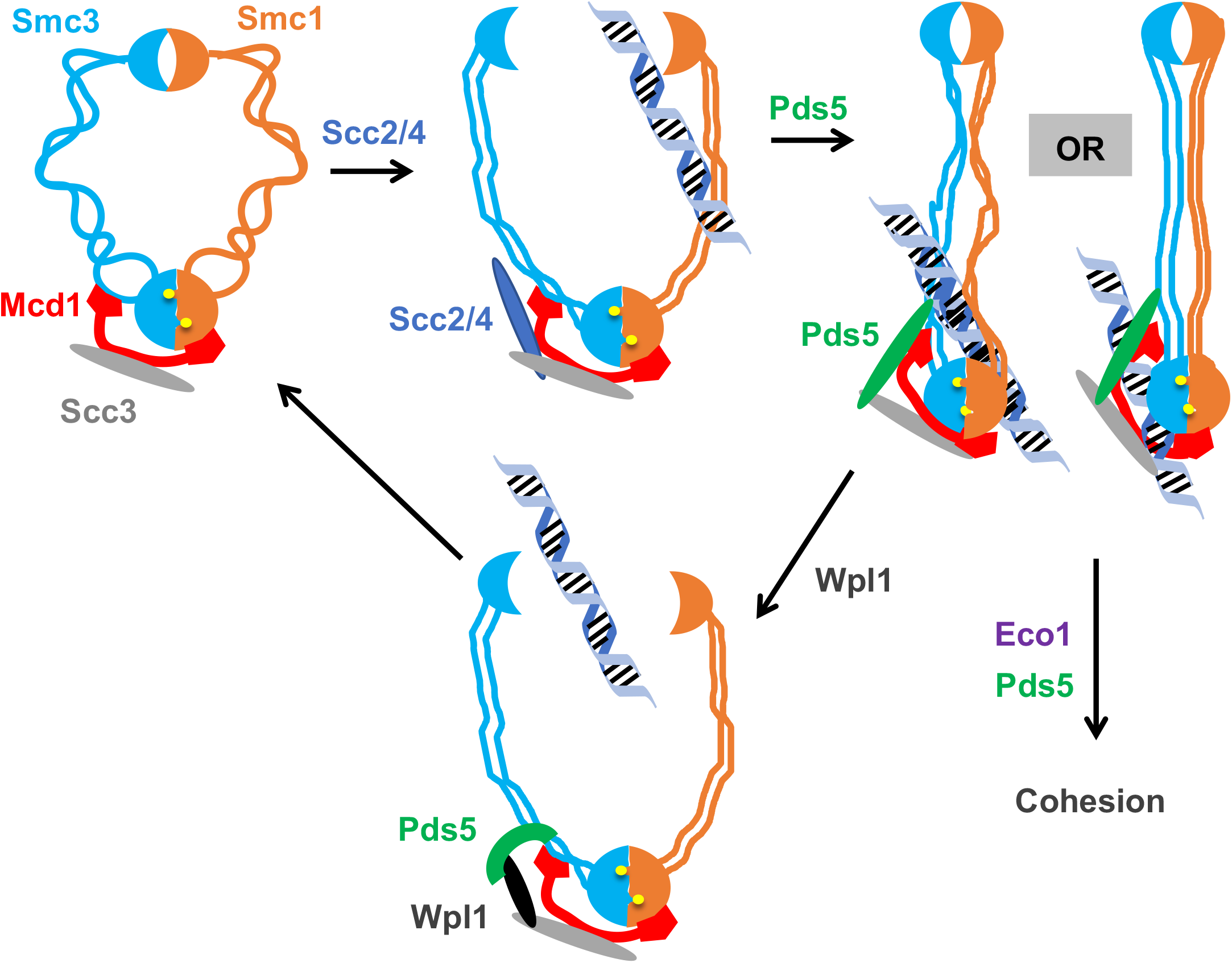
Model for how the Smc3p Mcd1p interface regulates the hinge dimer interface to control cohesin DNA. <u>Left side:</u> cohesin complex in an open ring conformation. <u>Middle top:</u> cohesin loader Scc2p/Scc4p (blue oval) binding at the Smc3p Mcd1p interface and triggering a conformation change that opens the hinge dimer (half-moons) interface, which allows DNA (blue-black helix) to enters cohesin. Alternatively, this conformational change could generate a folded ring to bring the hinges into close proximity to Scc2/Scc4 bound near the head, thereby enabling it to directly act on the hinges. <u>Right side:</u> DNA interaction with cohesin and Pds5p (green oval) binding at Smc3p Mcd1p interface triggers another conformational change that traps DNA either near the head (left) or in the small lumen formed by Mcd1p binding both the Smc1p head and the Smc3p coiled-coil (right). <u>Middle bottom:</u> Wpl1p binds Pds5p (green arc) and cohesin at its Smc3p Mcd1p interface and triggers a conformation change that opens hinge dimer (half-moons) interface and allows DNA to escape. Pds5p has both positive and negative functions so it is depicted differently with the green oval showing its positive role in promoting DNA binding and the green arc showing its negative role as it binds Wpl1p.

In summary, all SMC complexes have a conserved interface analogous to the interface between the N terminus of Mcd1p and the coiled coil of Smc3p (Gligoris and Löwe, 2016). Interestingly, a bacterial Smc complex and the yeast condensin can bind and then rapidly translocate along DNA (Terakawa *et al.*, 2017). These two activities will likely require conformational changes in the coiled coils. Given this conservation of the interfaces, these conformational changes are likely to be controlled by regulating novel functions of the Smc3p Mcd1p interface reported here.

## MATERIALS AND METHODS

### Yeast strains and media

Yeast strains used in this study are A364A background, and their genotypes are listed in Table 1. SC minimal and YPD media were prepared as described(Guacci *et al.*, 1997). Auxin (3-indole acetic acid; Sigma-Aldrich Catalog# I3705) Benomyl (a gift from Dupont) and camptothecin (Sigma catalog# C9911) plates used to assess drug sensitivity were prepared as previously described (Guacci and Koshland, 2012). Preparation of auxin containing media for depletion of AID tagged proteins was as previously described (Eng *et al.*, 2014).

### Dilution plating assays

Cells were grown to saturation in YPD media at 23°C (or 30°C when listed) then plated in 10-fold serial dilutions. Cells were incubated on plates at relevant temperatures or containing drugs as described.

### Synchronous arrest in mid-M phase under auxin depletion conditions

#### G1 arrest

Asynchronous cultures of cells were grown to mid-log phase at 23°C in YPD media, then alpha factor (αFactor) (Sigma; Catalog# T6901) was added to 10^-8^M. Cells were incubated for 23°C for 3h (or at 30°C for 2.5h) to induce arrest in G1 phase. For depletion of AID tagged proteins, auxin was added (500μM final) and cells incubated an additional 1h in αFactor containing media.

#### Synchronous arrest in mid-M phase

G1 arrested cells were washed 3x in YPD containing 0.1 mg/ml Pronase E (Sigma), once in YPD, then resuspended in YPD containing nocodozale (Sigma) at 15ug/ml final. For depletion of AID tagged proteins, auxin was added (500μM final) in all wash media and in resuspension media containing nocodazole to ensure depletion at all times. Cells were incubated at 23°C for 3h (or 30°C for 2.5h) to arrest in mid-M phase.

### Protein Extracts and Western Blotting

#### Total protein extracts

Cell equivalents of 2 to 4 OD_600_ were washed in cold 1xPBS, pelleted in a microfuge at 10k for 1 min then quick frozen using liquid nitrogen and stored at – 80°C. Cell extracts were made as described in (Eng *et al.*, 2015) with the following modifications. Initial lysis in an eppendorf tube was in 200ul 20% Trichloroacetic acid (W/V) then 500ul 5% TCA added twice, and all liquid combined in a new eppendorf and treated as described. The final protein pellet was resuspended in 212ul 2x Laemmli buffer + 26ul 1M Tris buffer pH8.

#### Immunoprecipitation

Cell equivalents of 20 OD_600_ were washed in cold 1xPBS, pelleted in a microfuge at 10k for 1 min then quick frozen using liquid nitrogen and stored at −80°C. Cells were lysed and cleared extracts incubated with Mouse anti-MYC (9E10) antibodies (Roche) to immunoprecipitate MYC epitope tagged proteins as previously described (Bloom *et al.*, 2018). Total extracts for these experiments were taken from cleared lysates before addition of antibodies.

#### Western Blots

Protein extracts were loaded onto 8% SDS page gels, subjected to electrophoresis then transferred to PDVF membranes using standard laboratory techniques.

### Monitoring cohesion using LacO-GFP assay

Cohesion was monitored using the LacO-LacI system. Briefly, cells contained a GFP-LacI fusion and tandem LacO repeats were integrated at *LYS4*, located 470kb from *CEN4.* Cells were fixed and processed to allow the number of GFP signals in each cell to be scored and the percentage of cells with 2-GFP spots determined as previously described (Guacci and Koshland, 2012).

**Chromatin Immunoprecipitation (ChIP)** was performed as previously described (Robison *et al.*, 2018). Primers used for ChIP are shown in Table 2.

#### Microscopy

Images were acquired with a Zeiss Axioplan2 microscope (100X objective, NA=1.40) equipped with a Quantix CCD camera (Photometrics).

**Flow cytometry analysis** was performed as previously described (Bloom *et al.*, 2018)..

### Plasmid constructs

Site directed mutagenesis using the Stratagene Quick-change kit was employed to generate the *smc3-I1026R* and *smc3-L1029R* alleles in normal cohesin on *LEU2* integrating plasmids pVG419 or in fusion cohesin on *LEU2* or *URA3* integrating plasmids pVG511 or p4897, respectively. Similarly, we generated *mcd1-L75K* and *mcd1-L75K* alleles in normal cohesin on *LEU2* integrating plasmids pJH18 or in fusion cohesin on *LEU2* integrating plasmids pVG511 respectively. The *smc3-K112R, K113R* mutant in fusion cohesin on *URA3* integrating plasmid p4897. The *smc3* and *mcd1* mutations were confirmed by sequencing the entire ORF and the promoter region to ensure it was the only change.

### Strain construction

#### *AID* tagged proteins

Details about the Auxin mediated destruction of AID tagged proteins in yeast was previously described (Eng *et al.*, 2014). Briefly, the *TIR1* E3-ubiquiting ligase placed under control of the GPD promoter and marked by *C. glibrata TRP1* replaced the *TRP1* gene on chromosome IV. Strains bearing genomic copies AID tagged of alleles of *SCC2, SCC3* and *PDS5* as sole source were made by C-terminal tag with *3V5-AID2* sequences using standard PCR techniques into TIR1 bearing yeast to generate *SCC2-AID, SCC3-AID* and *PDS5-AID* strains, respectively. *MCD1-AID* alleles were built the same way except using AID1 and did not have the 3V5 tag (Eng *et al.*, 2014). The *SMC3-AID* tagged strains were built as previously described (Guacci *et al.*, 2015).

## ACKNOWLEDGEMENTS

We thank Rebecca Lamothe, Lorenzo Constantino and Siheng Xiang for critical reading of the manuscript. This work was funded by National Institutes of Health Grant (1R35 GM-118189-01 to DK).

## COMPETING INTERESTS

The authors have no financial or non-financial competing interests.

---

**Supplemental Figure 1:**
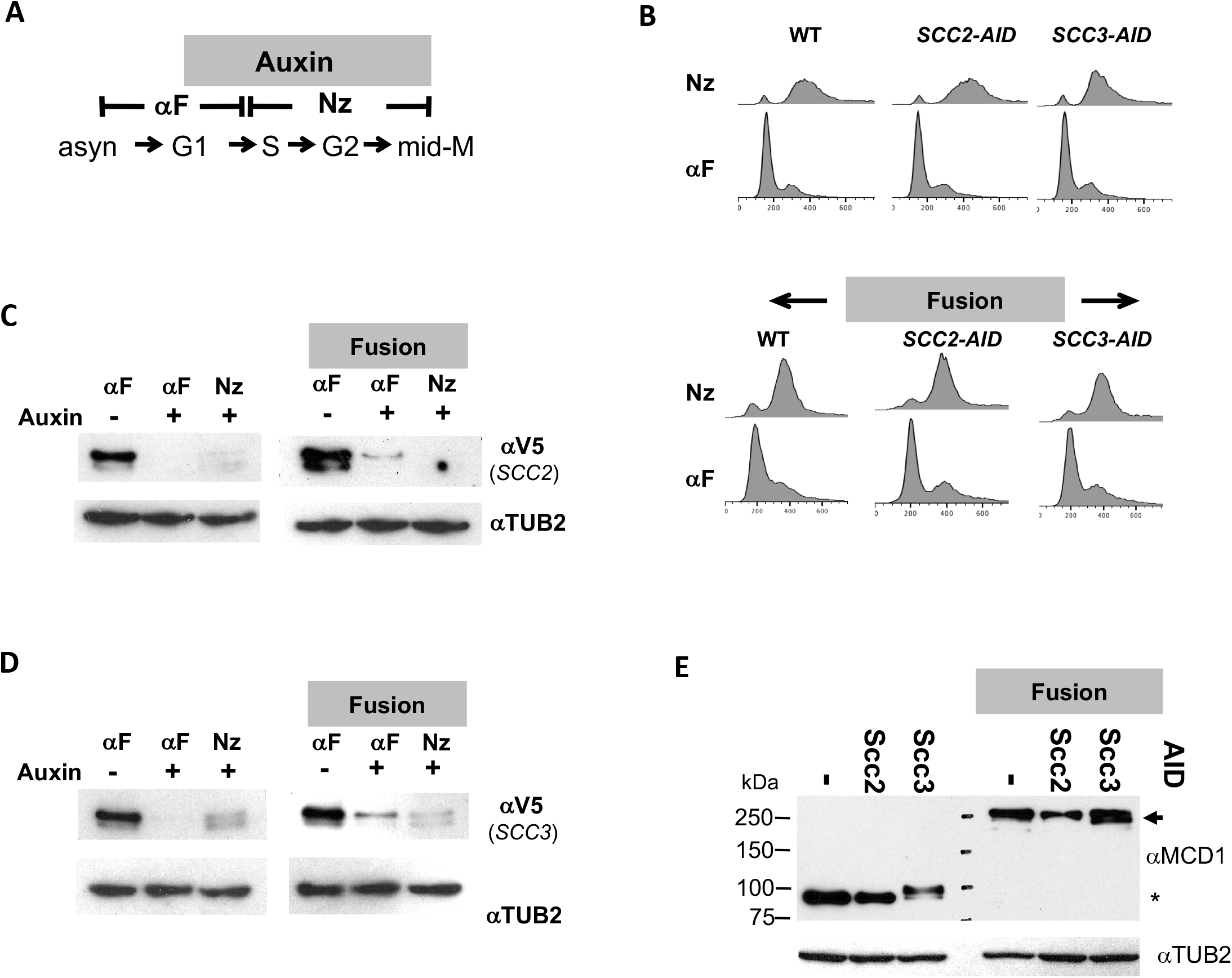
The effect of Scc2p-AID or Scc3p-AID depletion in normal and fusion cohesin strains. (A). Schematic of regimen used to synchronously arrest cells in mid-M phase. (B-D) Strains with normal cohesin or fusion cohesin alone or bearing Scc2p-AID or Scc3p-AID from Figure 2A were synchronously arrested in mid-M then processed to generate data presented in Figure 3A and 4A. (B) FACS to confirm arrest of cells. (C-D) Protein extracts were made from G1 arrested cells before and after auxin treatment, from synchronous arrest in mid-M were subjected to SDS-PAGE and analyzed by Western Blot. 3V5-AID tagged protein depletion was monitored using mouse antibodies against V5 (αV5) and rabbit anti-tubulin for a loading control (αTUB2). (C) Scc2p-3V5-AID depletion. (D) Scc3p-3V5-AID depletion (E) Mcd1p and Smc3p-Mcd1p fusion levels in mid-M arrested cells. Protein extracts from mid-M phase cells in C & D were analyzed by Western Blot. Mcd1p and fusion Smc3-Mcd1p fusion protein levels were monitored using rabbit antibodies against Mcd1p (αMCD1) and rabbit anti-tubulin for a loading control (αTUB2).

**Supplemental Figure 2:**
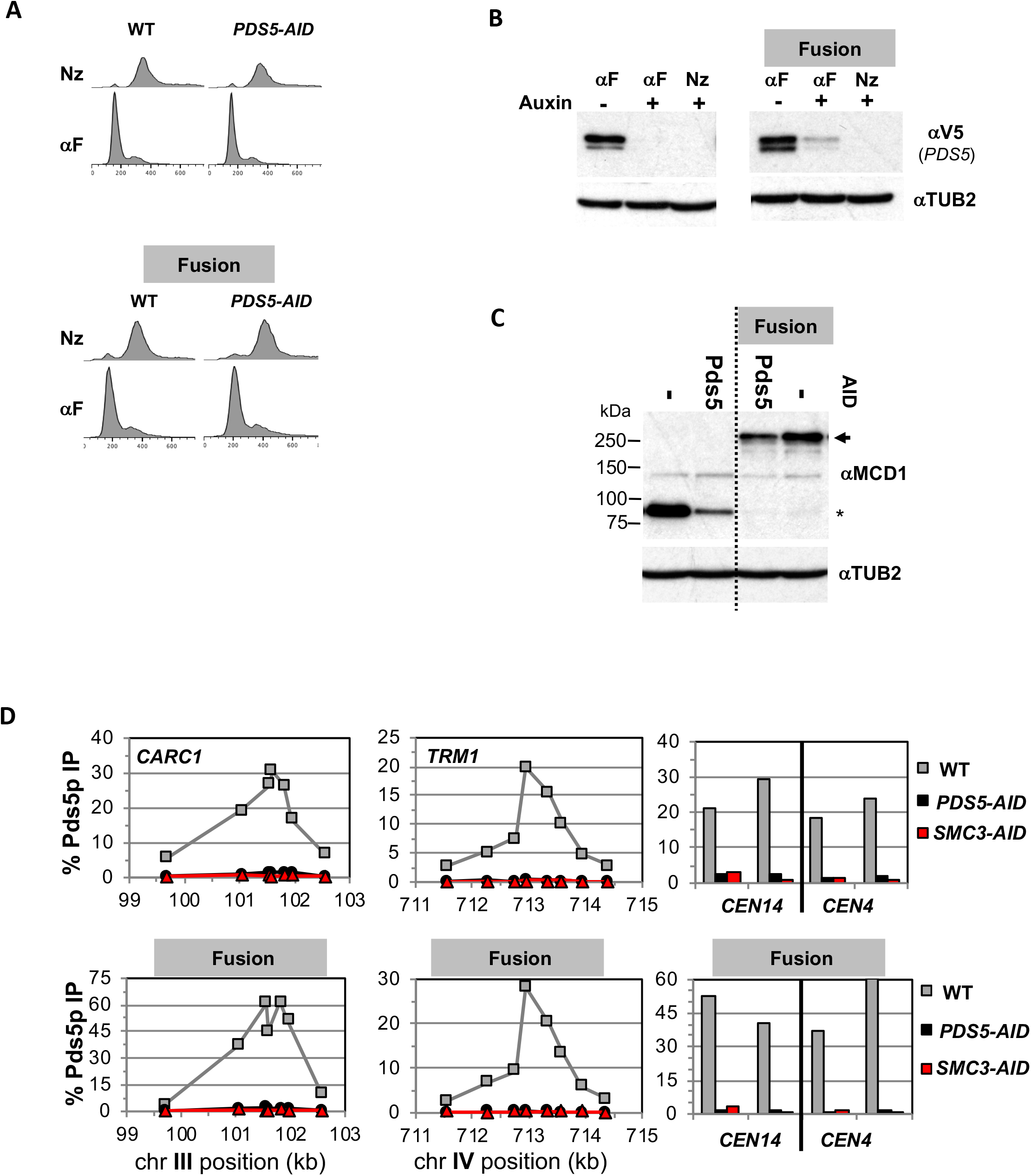
Comparing effects of Pds5p-AID depletion in normal and fusion cohesin strains. Strains with normal cohesin or fusion cohesin alone or bearing Pds5p-AID from Figure 2B were synchronously arrested in mid-M then processed to generate data presented in described in Figures 3B & 4B. (B) FACS to confirm arrest of cells. (B-C) Protein extracts were made from G1 arrested cells before and after auxin treatment and from synchronous arrest in mid-M were subjected to SDS-PAGE and analyzed by Western Blot. (B) Pds5p-3V5-AID depletion. 3V5-AID tagged protein depletion was monitored using mouse antibodies against V5 (αV5) and rabbit anti-tubulin for a loading control (αTUB2). depletion. (C) Mcd1p and Smc3p-Mcd1p fusion levels in mid-M arrested cells. Protein extracts from mid-M phase cells in B were analyzed by Western Blot. Mcd1p and Smc3-Mcd1p fusion protein levels were monitored using rabbit anti-Mcd1p antibodies (αMCD1) and rabbit anti-tubulin for a loading control (αTUB2). (D) Pds5p binding to chromosomes using ChIP. Mid-M phase cells from B-C were fixed and processed for ChIP to assess Pds5p binding after Pds5p depletion as described in Figure 3. Pds5p binding was assayed using rabbit anti-Pds5p antibodies (αPDS5). WT (grey), *PDS5-AID* (black) and *SMC3-AID* (red)

**Supplemental Figure 3:**
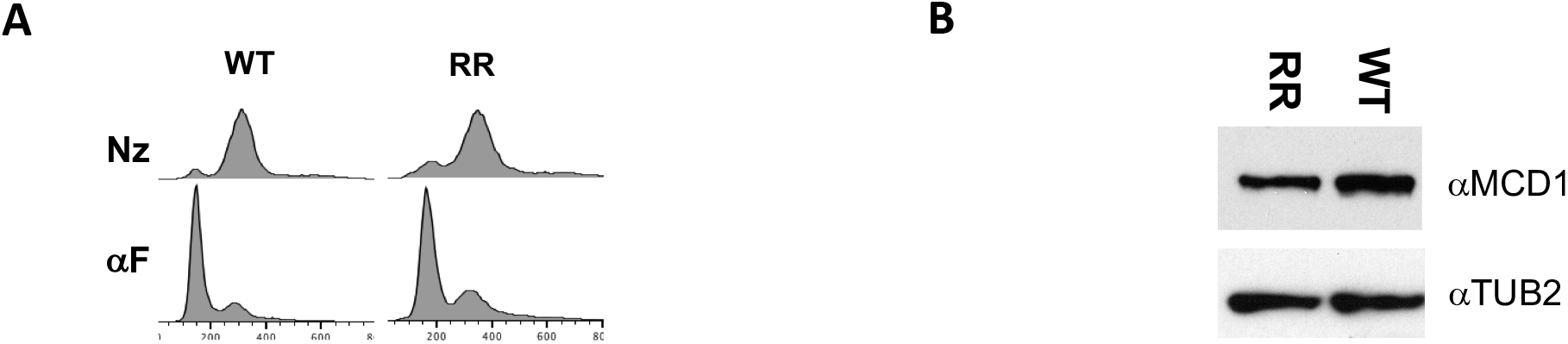
Characterization of fusion wild-type and *smc3-K112R K113R* (RR) mutations. Strains with fusion cohesin or RR fusion cohesin in figure 1C were grown treated as described in Figure 2C then processed to generate data presented in Figure 3C and 4C. (A) FACS to confirm arrest of cells. (B) Smc3p-Mcd1p fusion levels in mid-M arrested cells. Protein extracts from mid-M phase cells were analyzed by Western Blot. Smc3-Mcd1p fusion protein levels were monitored using rabbit anti-Mcd1p antibodies (αMCD1) and rabbit anti-tubulin for a loading control (αTUB2).

**Supplemental Figure 4:**
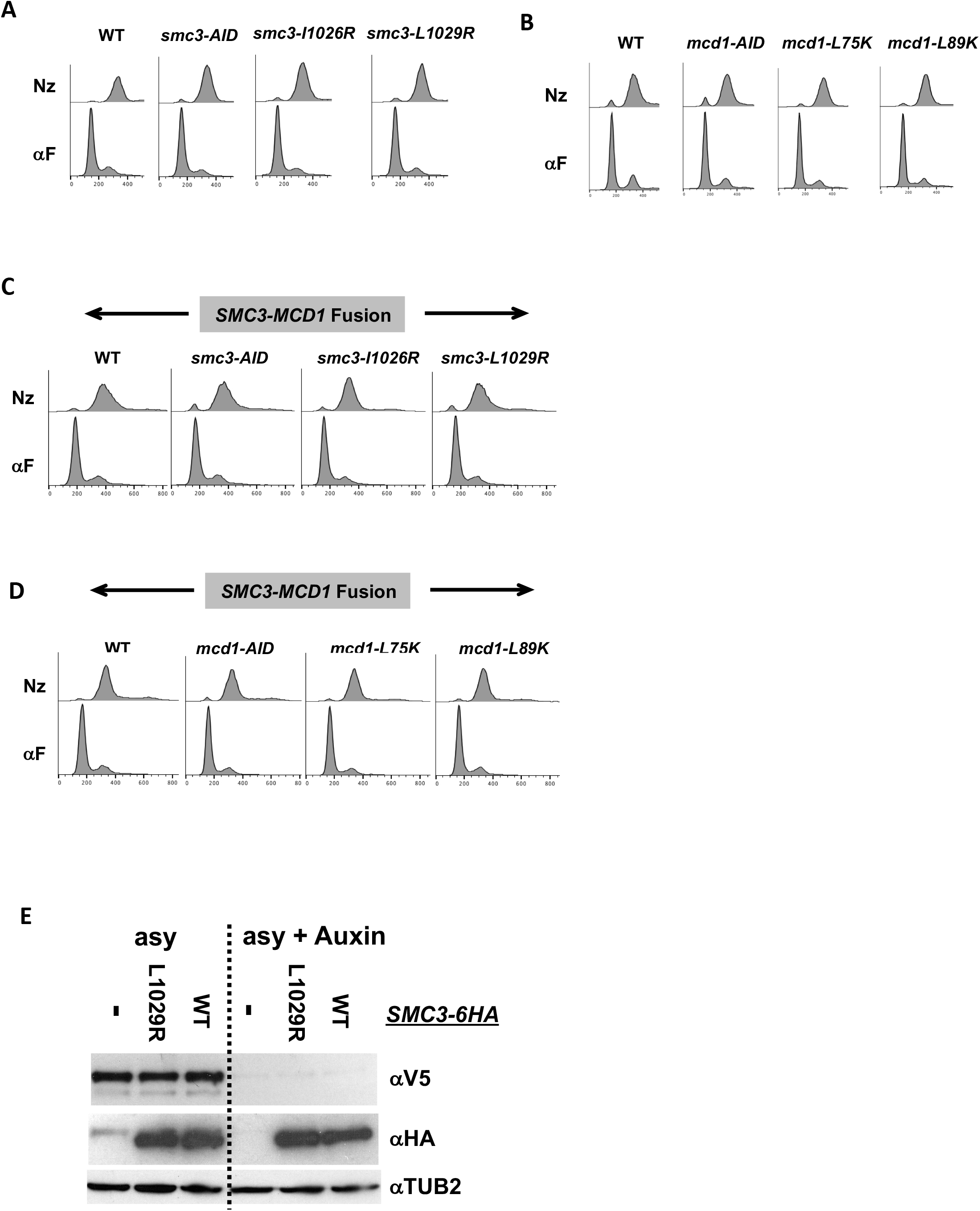
Characterization of normal and fusion cohesin bearing interface mutants. Strains with normal cohesin or fusion cohesin alone or bearing interface mutants from Figures 5 and 6 synchronously arrested in mid-M. (A-D) FACS analysis to confirm arrest. (A) *smc3* interface mutants in normal cohesin strains (B) *mcd1* interface mutants in normal cohesin strains. (C) *smc3* interface mutants in fusion cohesin strains. (D) *mcd1* interface mutants in fusion cohesin strains. (F) *smc3-L1029R* mutation has no effect on Smc3p protein level. Haploid normal cohesin strains bearing Smc3p-3V5-AID alone (VG3651-3D) or containing either Smc3p-6HA (VG3943-1C) or Smc3p-6HA-L1029R (VG3944-3D) were grown asynchronously, auxin added and cells incubated 1h. Protein extracts were made before after auxin addition and analyzed by Western Blot. Smc3p-6HA tagged proteins were monitored using mouse anti-HA antibodies (αHA), Smc3p-3V5-AID depletion was using mouse anti-V5 antibodies (αV5) and rabbit anti-tubulin for a loading control (αTUB2).

**Supplemental Figure 5:**
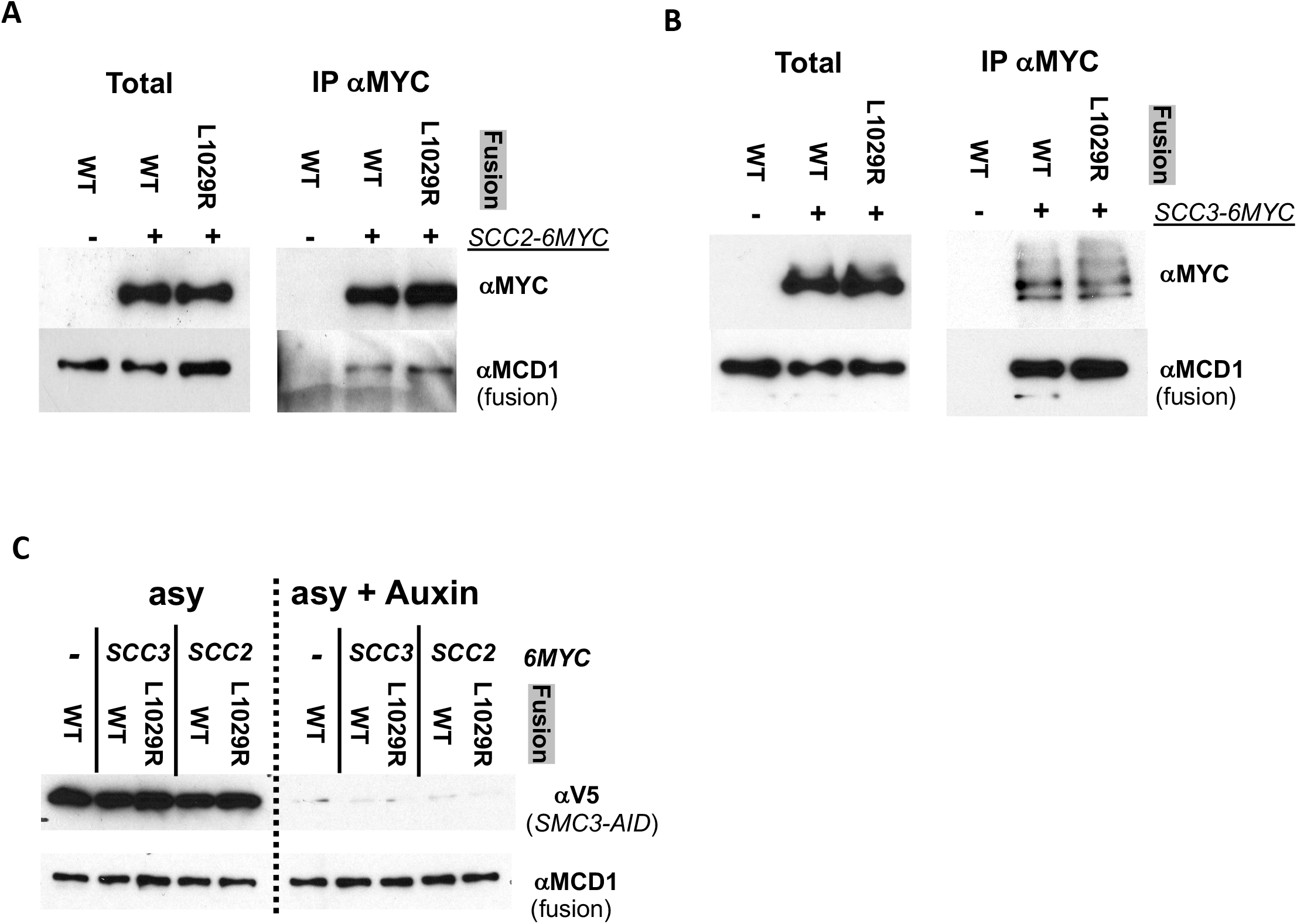
Fusion wild-type and fusion L1029 mutant coimmunoprecipitate with Scc2p and Scc3p equally well. (A) Fusion wild-type and L1029R mutant cohesin both co-IP with Scc2p. Haploid strain Scc2p-6MYC, Smc3p-3V5-AID bearing wild-type fusion cohesin (VG3952-14C) or fusion L1029R mutant (VG3953-17D) were grown asynchronously then auxin added and cells incubated 1h. An co-IP control without any MYC tagged proteins (VG3694-7C; wild-type fusion and Smc3p-3V5-AID) was also used. Protein extracts were made before after auxin addition. Mouse anti-MYC antibodies were used to immunoprecipitate MYC tagged proteins and analyzed by Western blot. Mouse anti-MYC antibodies detected MYC tagged Scc2p (αMYC), rabbit anti-Mcd1p antibodies detected fusion cohesin co-IP detected (αMCD1). (B) Fusion wild-type and L1029R mutant cohesin both co-IP with Scc3p. Haploid strain Scc3p-6MYC, Smc3p-3V5-AID bearing wild-type fusion cohesin (VG3949-5C) or fusion L1029R mutant (VG3950-8D) as well as control without any MYC tagged proteins (VG3694-7C; wild-type fusion and Smc3p-3V5-AID) were grown and treated as described in A. (C) Smc3p-3V5 depletion to confirm that fusion cohesin is sole cohesion in cells. Extracts from cells in A & B before and after auxin addition were analyzed by Western Blot. Smc3p-3V5-AID depletion was monitored using mouse anti-V5 antibodies (αV5) and fusion cohesin levels monitored using rabbit anti-Mcd1p antibodies (αMCD1).

## Yeast strains

VG3620-4C: *Mat****a*** *TIR1-CgTRP1 LacO-NAT::lys4 GFPLacI-HIS3:his3-11,15 leu2-3,112 ura3-52 bar1*
VG3630-7A: *Mat****a*** *G418:SCC2-3V5-AID2 TIR1-CgTRP1 LacO-NAT::lys4 GFPLacI-HIS3:his3-11,15 ura3-52 leu2-3,112 bar1*
VG3808-1A: *Mat****a*** *G418:SCC3-3V5-AID2 TIR1-CgTRP1 LacO-NAT::lys4 GFPLacI-HIS3:his3-11,15 ura3-52 leu2-3,112 bar1*
VG3940-2D: *Mat****a*** *SMC3-MCD1::*Δ*mcd1 smc3*Δ::*HPH LacO-NAT::lys4 GFPLacI-HIS3:his3-11,15 leu2-3,112 ura3-52 bar1*
VG3945-1A: *Mat****a*** *G418:SCC2-3V5-AID2 SMC3-MCD1::*Δ*mcd1 smc3*Δ::*HPH LacO-NAT::lys4 GFPLacI-HIS3:his3-11,15 leu2-3,112 ura3-52 bar1*
VG3946-7B: *Mat****a*** *G418:SCC3-3V5-AID2 SMC3-MCD1::*Δ*mcd1 smc3*Δ::*HPH LacO-NAT::lys4 GFPLacI-HIS3:his3-11,15 leu2-3,112 ura3-52 bar1*
VG3954-10C: *Mat****a*** *G418:PDS5-3V5-AID2 TIR1-CgTRP1 LacO-NAT::lys4 GFPLacI-HIS3:his3-11,15 leu2-3,112 ura3-52 bar1*
VG3955-4D: *Mat****a*** *G418:PDS5-3V5-AID2 SMC3-MCD1::*Δ*mcd1 smc3*Δ::*HPH LacO-NAT::lys4 GFPLacI-HIS3:his3-11,15 leu2-3,112 ura3-52 bar1*
VG3930-5C: *Mat****a*** *SMC3-K112R,K113R-MCD1::*Δ*mcd1 smc3*Δ::*HPH LacO-NAT::lys4 leu2-3,112 GFPLacI-HIS3:his3-11,15 ura3-52 bar1*
VG3651-3D: *Mat****a*** *SMC3-3V5-AID*^*608*^ *TIR1-CgTRP1 LacO-NAT::lys4 GFPLacI-HIS3:his3-11,15 leu2-3,112 ura3-52 bar1*
BRY474: *Mat****a*** *SMC3-LEU2:leu2-3,112 SMC3-3V5-AID*^*608*^ *TIR1-CgTRP1 LacO-NAT::lys4 GFPLacI-HIS3:his3-11,15 ura3-52 bar1*
BRY492: *Mat****a*** *smc3-L1029R-LEU2:leu2-3,112 SMC3-3V5-AID*^*608*^ *TIR1-CgTRP1 LacO-NAT::lys4 GFPLacI-HIS3:his3-11,15 ura3-52 bar1*
VG3905-7A: *Mat****a*** *smc3-I1026R-LEU2:leu2-3,112 SMC3-3V5-AID*^*608*^ *TIR1-CgTRP1 LacO-NAT::lys4 GFPLacI-HIS3:his3-11,15 ura3-52 bar1*
VG3943-1C: *Mat****a*** *SMC3-6HA*^*F864*^*-LEU2:leu2-3,112 SMC3-3V5-AID*^608^ *TIR1-CgTRP1 LacO-NAT::lys4 GFPLacI-HIS3:his3-11,15 ura3-52 bar1*
VG3944-3D: *Mat****a*** *smc3-6HA*^*F864*^*-I1029R-LEU2:leu2-3,112 SMC3-3V5-AID*^608^ *TIR1-CgTRP1 LacO-NAT::lys4 GFPLacI-HIS3:his3-11,15 ura3-52 bar1*
VG3902-3A: *Mat****a*** *G418:MCD1-AID TIR1-CgTRP1 LacO-NAT::lys4 GFPLacI-HIS3:his3-11,15 leu2-3,112 ura3-52 bar1*
VG3914-1C: *Mat****a*** *MCD1-LEU2:leu2-3,112 G418:MCD1-AID TIR1-CgTRP1 LacO-NAT::lys4 GFPLacI-HIS3:his3-11,15 leu2-3,112 ura3-52 bar1*
VG3916-5B: *Mat****a*** *mcd1-L75K-LEU2:leu2-3,112 G418:MCD1-AID TIR1-CgTRP1 LacO-NAT::lys4 GFPLacI-HIS3:his3-11,15 leu2-3,112 ura3-52 bar1*
VG3918-9D: *Mat****a*** *mcd1-L89K-LEU2:leu2-3,112 G418:MCD1-AID TIR1-CgTRP1 LacO-NAT::lys4 GFPLacI-HIS3:his3-11,15 leu2-3,112 ura3-52 bar1*
VG3694-7C: *Mat****a*** *SMC3MCD1-URA3:ura3-52 SMC3-3V5-AID*^*608*^ *TIR1-CgTRP1 LacO-NAT::lys4 GFPLacI-HIS3:his3-11,15 leu2-3,112 bar1*
VG3908-17B: *Mat****a*** *smc3-I1026R-MCD1-URA3:ura3-52 SMC3-3V5-AID*^*608*^ *TIR1-CgTRP1 LacO-NAT::lys4 GFPLacI-HIS3:his3-11,15 leu2-3,112 bar1*
VG3872-3B: *Mat****a*** *smc3-L1029R-MCD1-URA3:ura3-52 SMC3-3V5-AID*^*608*^ *TIR1-CgTRP1 LacO-NAT::lys4 GFPLacI-HIS3:his3-11,15 leu2-3,112 bar1*
VG3937-2C: *Mat****a*** *SMC3MCD1-LEU2:leu2-3,112 G418:MCD1-AID TIR1-CgTRP1 LacO-NAT::lys4 GFPLacI-HIS3:his3-11,15 ura3-52 bar1*
VG3938-3A: *Mat****a*** *SMC3mcd1-L75K-LEU2:leu2-3,112 G418:MCD1-AID TIR1-CgTRP1 LacO-NAT::lys4 GFPLacI-HIS3:his3-11,15 ura3-52 bar1*
VG3939-7B: *Mat****a*** *SMC3mcd1-L89K-LEU2:leu2-3,112 G418:MCD1-AID TIR1-CgTRP1 LacO-NAT::lys4 GFPLacI-HIS3:his3-11,15 ura3-52 bar1*
VG3952-14C: *Mat****a*** *G418:SCC2-6MYC SMC3MCD1-URA3:ura3-52 SMC3-3V5-AID*^608^ *TIR1-CgTRP1 LacO-NAT::lys4 GFPLacI-HIS3:his3-11,15 leu2-3,112 bar1*
VG3953-17D: *Mat****a*** *G418:SCC2-6MYC smc3-L1029R-MCD1-URA3:ura3-52 SMC3-3V5-AID*^608^ *TIR1-CgTRP1 LacO-NAT::lys4 GFPLacI-HIS3:his3-11,15 leu2-3,112 bar1*
VG3949-5C: *Mat****a*** *G418:SCC3-6MYC SMC3-MCD1-URA3:ura3-52 SMC3-3V5-AID*^608^ *TIR1-CgTRP1 LacO-NAT::lys4 GFPLacI-HIS3:his3-11,15 leu2-3,112 bar1*
VG3950-8D: *Mat****a*** *G418:SCC3-6MYC smc3-L1029R-MCD1-URA3:ura3-52 SMC3-3V5-AID*^608^ *TIR1-CgTRP1 LacO-NAT::lys4 GFPLacI-HIS3:his3-11,15 leu2-3,112 bar1*
VG3927-14D: *Mat****a*** *MCD1-S83D TIR1-CgTRP1 LacO-NAT::lys4 GFPLacI-HIS3:his3-11,15 leu2-3,112 ura3-52 bar1*

## Primers used for chromatin Immunoprecipitation (ChIP

### CEN-distal TRM1 CAR

<u>VG641/VG642</u> (5’ AAA GAA GCA GGG GTA GAG AAG C 3’/ 5’ ATC AGC AGC GGT GAT TAC AC 3’)

<u>VG625/VG626</u> (5’ CCA GCC AGA TAT TAT GGG CAA G 3’/ 5’ TCC TAG ACC TGG TGG AAA AAG C 3’)

<u>VG627/VG628</u> (5’ CTT ATA GTT CCC AAG GCA TCC C 3’/ 5’ CCA AAC TCG TTG TTC TCG ATC C 3’)

<u>VG629/VG630</u> (5’ TCT TCG TGC GCG AGG ATA TG 3’/ 5’ CGA ACA TTT CCG GAC AAT TGC 3’)

<u>VG633/VG634</u> (5’ CCA ATC GTA TAA CGG AGC ATT GG 3’/ 5’ TGG TGC CAG AAG ATA TCA ACG 3’)

<u>VG637/VG638</u> (5’ GCG CGA TAC CAT TCA GAA CAT C 3’/ 5’ TTA AAG TGG GCC CAA GAC CAG 3’)

<u>VG639/VG640</u> (5’ GGG CAT CAC CTT TTC GTA AGC 3’ / 5’ TGA TCC ACC TGT CAT TTC GC 3’)

<u>VG643/VG644</u> (5’ ACC CTT CTG TTC CAG TTT GC 3’/ 5’ GTT GCC TCC GGA GCA AAT TC 3’)

### Pericentric CARC1

<u>TE367/TE368</u> (5’ AAA GGT GCC CCA AGA AAA GG 3’/ 5’ AGC ACT TTA CTC GCT TGT GG 3’)

<u>TE310/TE311</u> (5’ TAA AGC ATT GAC GCC AGA GC3’/ 5’ AAG TAC GCG TAC GAA GCA TC 3’)

<u>TE306/TE307</u> (5’ CAA ACC ACC TCT TAC GTC GTT G3’/ 5’ TTT CGT GCA CTG CGT TCA AG 3’)

<u>TE308/TE309</u> (5’ TCC TGG AAT GGA GAC CGT TTT C 3’/ 5’ AGC CGA CAA ATT TCG TGC AC 3’)

<u>TE373/TE374</u> (5’ ACT TTG GTT TTC CGG TGT GC3’/ 5’ CCA GCG ATG AGA TGC GAA AAG 3’)

<u>TE377/TE378</u> (5’ TCG CTT TTC GCA TCT CAT CG 3’/ 5’ AGC GGG CGG GTT ATA AAT AAC 3’)

<u>TE533/TE534</u> (5’ ACC TTC TAC TTC CAT GCC GTT G 3’/ 5’ TGC GTG CCG ATG TAG AAT TG 3’)

### CEN primers

#### CEN4 flanking

<u>BR463/BR464</u> (5’ CAT GAT TCG CCG GGT AAA TA 3’/ 5’ GCA CTA GCC AAT TTA GCA CTT C 3’)

<u>BR465/BR466</u> (5’ AAA ATG CCG AGG CTT TCA TA 3’/ 5’ TGA CGA TAA AAC CGG AAG GA 3’

#### CEN14 flanking

<u>TE442/TE443</u> (5’ TTA AAG CGG CTG AGT ATG GC 3’/ 5’ TTT CCT CCA TTG CTC TCT ACG G 3’)

<u>TE446/TE447</u> (5’ ACT AAA AGT GCC CCA AAC GG 3’/ 5’ AGG AGC AGG GTA GCA TAA ACC 3’)

## REFERENCES

Arumugam, P., Nishino, T., Haering, C. H., Gruber, S., and Nasmyth, K. (2006). Cohesin’s ATPase activity is stimulated by the C-terminal Winged-Helix domain of its kleisin subunit. Curr Biol 16, 1998–2008.

Beckouet, F. et al. (2016). Releasing Activity Disengages Cohesin’s Smc3/Scc1 Interface in a Process Blocked by Acetylation. Mol Cell 61, 563–574.

Blat, Y., and Kleckner, N. (1999). Cohesins bind to preferential sites along yeast chromosome III, with differential regulation along arms versus the centric region. Cell 98, 249–259.

Bloom, M. S., Koshland, D., and Guacci, V. (2018). Cohesin Function in Cohesion, Condensation, and DNA Repair Is Regulated by Wpl1p via a Common Mechanism in Saccharomyces cerevisiae. Genetics 208, 111–124.

Chan, K.-L., Gligoris, T., Upcher, W., Kato, Y., Shirahige, K., Nasmyth, K., and Beckouet, F. (2013). Pds5 promotes and protects cohesin acetylation. Proc Natl Acad Sci USA 110, 13020–13025.

Chan, K.-L., Roig, M. B., Hu, B., Beckouet, F., Metson, J., and Nasmyth, K. (2012). Cohesin’s DNA exit gate is distinct from its entrance gate and is regulated by acetylation. Cell 150, 961–974.

Ciosk, R., Shirayama, M., Tanaka, T., Toth, A., Shevchenko, A., and Nasmyth, K. (2000). Cohesin’s binding to chromosomes depends on a separate complex consisting of Scc2 and Scc4 proteins. Mol Cell 5, 243–254.

Çamdere, G. Ö., Carlborg, K. K., and Koshland, D. (2018). Intermediate step of cohesin’s ATPase cycle allows cohesin to entrap DNA. Proc Natl Acad Sci USA 115, 9732–9737.

Çamdere, G., Guacci, V., Stricklin, J., and Koshland, D. (2015). The ATPases of cohesin interface with regulators to modulate cohesin-mediated DNA tethering. Elife 4, 13115.

D’Ambrosio, L. M., and Lavoie, B. D. (2014). Pds5 Prevents the PolySUMO-Dependent Separation of Sister Chromatids. Current Biology 24, 361–371.

Eagen, K. P. (2018). Principles of Chromosome Architecture Revealed by Hi-C. Trends Biochem Sci 43, 469–478.

Elbatsh, A. M. O. et al. (2016). Cohesin Releases DNA through Asymmetric ATPase-Driven Ring Opening. Mol Cell 61, 575–588.

Eng, T., Guacci, V., and Koshland, D. (2014). ROCC, a conserved region in cohesin’s Mcd1 subunit, is essential for the proper regulation of the maintenance of cohesion and establishment of condensation. Mol Biol Cell 25, 2351–2364.

Eng, T., Guacci, V., and Koshland, D. (2015). Interallelic complementation provides functional evidence for cohesin-cohesin interactions on DNA. Mol Biol Cell 26, 4224–4235.

Gandhi, R., Gillespie, P. J., and Hirano, T. (2006). Human Wapl Is a Cohesin-Binding Protein that Promotes Sister-Chromatid Resolution in Mitotic Prophase. Current Biology 16, 2406–2417.

Gerlich, D., Koch, B., Dupeux, F., Peters, J.-M., and Ellenberg, J. (2006). Live-cell imaging reveals a stable cohesin-chromatin interaction after but not before DNA replication. Curr Biol 16, 1571–1578.

Gligoris, T. G., Scheinost, J. C., Burmann, F., Petela, N., Chan, K. L., Uluocak, P., Beckouet, F., Gruber, S., Nasmyth, K., and Lowe, J. (2014). Closing the cohesin ring: Structure and function of its Smc3-kleisin interface. Science 346, 963–967.

Gligoris, T., and Löwe, J. (2016). Structural Insights into Ring Formation of Cohesin and Related Smc Complexes. Trends Cell Biol. 26, 680–693.

Gruber, S., Arumugam, P., Katou, Y., Kuglitsch, D., Helmhart, W., Shirahige, K., and Nasmyth, K. (2006). Evidence that Loading of Cohesin Onto Chromosomes Involves Opening of Its SMC Hinge. Cell 127, 523–537.

Guacci, V., and Koshland, D. (2012). Cohesin-independent segregation of sister chromatids in budding yeast. Mol Biol Cell 23, 729–739.

Guacci, V., Hogan, E., and Koshland, D. (1994). Chromosome condensation and sister chromatid pairing in budding yeast. J Cell Biol 125, 517–530.

Guacci, V., Koshland, D., and Strunnikov, A. (1997). A direct link between sister chromatid cohesion and chromosome condensation revealed through the analysis of MCD1 in S. cerevisiae. Cell 91, 47–57.

Guacci, V., Stricklin, J., Bloom, M. S., Guō, X., Bhatter, M., and Koshland, D. (2015). A novel mechanism for the establishment of sister chromatid cohesion by the ECO1 acetyltransferase. Mol Biol Cell 26, 117–133.

Haering, C. H., Farcas, A.-M., Arumugam, P., Metson, J., and Nasmyth, K. (2008). The cohesin ring concatenates sister DNA molecules. Nature 454, 297–301.

Hartman, T., Stead, K., Koshland, D., and Guacci, V. (2000). Pds5p is an essential chromosomal protein required for both sister chromatid cohesion and condensation in Saccharomyces cerevisiae. J Cell Biol 151, 613–626.

Heidinger-Pauli, J. M., Unal, E., and Koshland, D. (2009). Distinct targets of the Eco1 acetyltransferase modulate cohesion in S phase and in response to DNA damage. Mol Cell 34, 311–321.

Huber, R. G. et al. (2016). Impairing Cohesin Smc1/3 Head Engagement Compensates for the Lack of Eco1 Function. Structure 24, 1991–1999.

Huis in ’t Veld, P. J., Herzog, F., Ladurner, R., Davidson, I. F., Piric, S., Kreidl, E., Bhaskara, V., Aebersold, R., and Peters, J.-M. (2014). Characterization of a DNA exit gate in the human cohesin ring. Science 346, 968–972.

Ivanov, D., and Nasmyth, K. (2007). A physical assay for sister chromatid cohesion in vitro. Mol Cell 27, 300–310.

Kueng, S., Hegemann, B., Peters, B. H., Lipp, J. J., Schleiffer, A., Mechtler, K., and Peters, J.-M. (2006). Wapl controls the dynamic association of cohesin with chromatin. Cell 127, 955–967.

Laloraya, S., Guacci, V., and Koshland, D. (2000). Chromosomal addresses of the cohesin component Mcd1p. J Cell Biol 151, 1047–1056.

Lavoie, B. D., Hogan, E., and Koshland, D. (2002). In vivo dissection of the chromosome condensation machinery: reversibility of condensation distinguishes contributions of condensin and cohesin. J Cell Biol 156, 805–815.

Lengronne, A., McIntyre, J., Katou, Y., Kanoh, Y., Hopfner, K.-P., Shirahige, K., and Uhlmann, F. (2006). Establishment of Sister Chromatid Cohesion at the S. cerevisiae Replication Fork. Mol Cell 23, 787–799.

Lopez-Serra, L., Lengronne, A., Borges, V., Kelly, G., and Uhlmann, F. (2013). Budding Yeast Wapl Controls Sister Chromatid Cohesion Maintenance and Chromosome Condensation. Current Biology 23, 64–69.

Losada, A., Hirano, M., and Hirano, T. (1998). Identification of Xenopus SMC protein complexes required for sister chromatid cohesion. Genes Dev 12, 1986–1997.

Megee, P. C., Mistrot, C., Guacci, V., and Koshland, D. (1999). The centromeric sister chromatid cohesion site directs Mcd1p binding to adjacent sequences. Mol Cell 4, 445–450.

Michaelis, C., Ciosk, R., and Nasmyth, K. (1997). Cohesins: chromosomal proteins that prevent premature separation of sister chromatids. Cell.

Murayama, Y., and Uhlmann, F. (2013). Biochemical reconstitution of topological DNA binding by the cohesin ring. Nature 505, 367–371.

Murayama, Y., and Uhlmann, F. (2015). DNA Entry into and Exit out of the Cohesin Ring by an Interlocking Gate Mechanism. Cell 163, 1628–1640.

Noble, D., Kenna, M. A., Dix, M., Skibbens, R. V., Unal, E., and Guacci, V. (2006). Intersection between the regulators of sister chromatid cohesion establishment and maintenance in budding yeast indicates a multi-step mechanism. Cell Cycle 5, 2528–2536.

Onn, I., Heidinger-Pauli, J. M., Guacci, V., Unal, E., and Koshland, D. E. (2008). Sister chromatid cohesion: a simple concept with a complex reality. 24, 105–129.

Orgil, O., Matityahu, A., Eng, T., Guacci, V., Koshland, D., and Onn, I. (2015). A Conserved Domain in the Scc3 Subunit of Cohesin Mediates the Interaction with Both Mcd1 and the Cohesin Loader Complex. PLoS Genet 11, e1005036.

Robison, B., Guacci, V., and Koshland, D. (2018). A role for the Smc3 hinge domain in the maintenance of sister chromatid cohesion. Mol Biol Cell 29, 339–355.

Roig, M. B., Löwe, J., Chan, K.-L., Beckouet, F., Metson, J., and Nasmyth, K. (2014). Structure and function of cohesin’s Scc3/SA regulatory subunit. FEBS Lett 588, 3692–3702.

Rolef Ben-Shahar, T., Heeger, S., Lehane, C., East, P., Flynn, H., Skehel, M., and Uhlmann, F. (2008). Eco1-dependent cohesin acetylation during establishment of sister chromatid cohesion. Science 321, 563–566.

Rollins, R. A., Korom, M., Aulner, N., Martens, A., and Dorsett, D. (2004). Drosophila nipped-B protein supports sister chromatid cohesion and opposes the stromalin/Scc3 cohesion factor to facilitate long-range activation of the cut gene. Mol Cell Biol 24, 3100–3111.

Rowland, B. D. et al. (2009). Building sister chromatid cohesion: smc3 acetylation counteracts an antiestablishment activity. Mol Cell 33, 763–774.

Skibbens, R. V., Corson, L. B., Koshland, D., and Hieter, P. (1999). Ctf7p is essential for sister chromatid cohesion and links mitotic chromosome structure to the DNA replication machinery. Genes Dev 13, 307–319.

Soh, Y.-M. et al. (2015). Molecular Basis for SMC Rod Formation and Its Dissolution upon DNA Binding. Mol Cell 57, 290–303.

Stead, K., Aguilar, C., Hartman, T., Drexel, M., Meluh, P., and Guacci, V. (2003). Pds5p regulates the maintenance of sister chromatid cohesion and is sumoylated to promote the dissolution of cohesion. J Cell Biol 163, 729–741.

Stigler, J., Çamdere, G. Ö., Koshland, D. E., and Greene, E. C. (2016). Single-Molecule Imaging Reveals a Collapsed Conformational State for DNA-Bound Cohesin. CellReports 15, 988–998.

Strom, L., Karlsson, C., Lindroos, H. B., Wedahl, S., Katou, Y., Shirahige, K., and Sjögren, C. (2007). Postreplicative formation of cohesion is required for repair and induced by a single DNA break. Science 317, 242–245.

Sutani, T., Kawaguchi, T., Kanno, R., Itoh, T., and Shirahige, K. (2009). Budding Yeast Wpl1(Rad61)-Pds5 Complex Counteracts Sister Chromatid Cohesion-Establishing Reaction. Current Biology, 1–6.

Terakawa, T., Bisht, S., Eeftens, J. M., Dekker, C., Haering, C. H., and Greene, E. C. (2017). The condensin complex is a mechanochemical motor that translocates along DNA. Science 358, 672–676.

Tóth, A., Ciosk, R., Uhlmann, F., Galova, M., Schleiffer, A., and Nasmyth, K. (1999). Yeast cohesin complex requires a conserved protein, Eco1p(Ctf7), to establish cohesion between sister chromatids during DNA replication. Genes Dev 13, 320–333.

Unal, E., Heidinger-Pauli, J. M., and Koshland, D. (2007). DNA double-strand breaks trigger genome-wide sister-chromatid cohesion through Eco1 (Ctf7). Science 317, 245–248.

Unal, E., Heidinger-Pauli, J. M., Kim, W., Guacci, V., Onn, I., Gygi, S. P., and Koshland, D. E. (2008). A molecular determinant for the establishment of sister chromatid cohesion. Science 321, 566–569.

